# The blast effector Pwl2 is a virulence factor that modifies the cellular localisation of host protein HIPP43 to suppress immunity

**DOI:** 10.1101/2024.01.20.576406

**Authors:** Vincent Were, Xia Yan, Andrew J. Foster, Jan Sklenar, Thorsten Langner, Amber Gentle, Neha Sahu, Adam Bentham, Rafał Zdrzałek, Lauren Ryder, Davies Kaimenyi, Diana Gomez De La Cruz, Yohan Petit-Houdenot, Alice Bisola Eseola, Matthew Smoker, Mark Jave Bautista, Weibin Ma, Jiorgos Kourelis, Dan Maclean, Mark J. Banfield, Sophien Kamoun, Frank L.H. Menke, Matthew J. Moscou, Nicholas J. Talbot

## Abstract

The rice blast fungus *Magnaporthe oryzae* secretes a battery of effector proteins to facilitate host infection. Among these effectors, Pwl2 was first identified as a host specificity determinant for infection of weeping lovegrass (*Eragrostis curvula*) and is also recognised by the barley Mla3 resistance gene. However, its biological activity is not known. Here we show that *PWL2* expression is regulated by the Pmk1 MAP kinase during cell-to-cell movement by *M. oryzae* at plasmodesmata (PD)-containing pit field sites. Consistent with its regulation, we provide evidence that Pwl2 binds to a barley heavy metal-binding isoprenylated protein HIPP43, which results in its displacement from plasmodesmata. Transgenic barley lines overexpressing either *PWL2* or HIPP43 exhibit attenuated immune responses and increased disease susceptibility. By contrast, a Pwl2^SNDEYWY^ mutant that does not interact with HIPP43, fails to alter the PD localisation of HIPP43. Targeted deletion of three copies of *PWL2* in *M. oryzae* results in a *Δpwl2* mutant showing gain-of-virulence to weeping lovegrass and barley Mla3 lines, but also a reduction in severity of blast disease on susceptible host plants. Taken together, our results provide evidence that Pwl2 is a virulence factor that acts by suppressing host immunity through perturbing the plasmodesmatal deployment of HIPP43.

## Introduction

Plant pathogens secrete virulence proteins called effectors during infection to suppress plant immunity and facilitate infection (Jones and Dangl, 2006). Fungal pathogens, such as the devastating blast fungus *Magnaporthe oryzae*, utilise an extensive battery of more than 500 effectors (Wilson and Talbot, 2009; Yan *et al*., 2023), but very few have been functionally characterised (Oliveira-Garcia *et al*., 2023a). A subset of effectors are recognized by plant intracellular nucleotide-binding leucine rich repeat (NLR) immune receptors to activate disease resistance. In rice for example, the paired NLRs RGA5/RGA4, Pik-1/Pik-2, and Piks-1/Piks-2 confer resistance to *M. oryzae* strains that secrete AVR1-CO39/AVR-Pia, AVR-Pik or AVR-Mgk1 effectors, respectively (Ashikawa *et al*., 2008; Kanzaki *et al*., 2012; Cesari *et al*., 2013; Ortiz *et al*., 2017; Zhang *et al*., 2018; Sugihara *et al*., 2023). These effectors are members of the *Magnaporthe* Avrs and ToxB-like (MAX) family which are sequence-unrelated, structurally conserved (de Guillen *et al*., 2015) and overrepresented among effectors recognised by rice NLRs (Cesari *et al*., 2013; Maqbool *et al*., 2015). Pwl2 is a MAX effector first identified as a host specificity determinant that controls pathogenicity towards weeping lovegrass (*Eragrostis curvula*), a widely-grown forage grass (Sweigard *et al*., 1995), and was recently shown to be recognised in barley by the Mla3 immune receptor (Brabham *et al*., 2023). While Pwl2 has been widely studied to understand effector secretion and delivery (Mosquera *et al*., 2009; Zhang and Xu, 2014; Oliveira-Garcia *et al*., 2023b), the biological function of Pwl2 is unknown.

Recent studies of MAX-effector perception by NLRs have implicated host small heavy metal-associated (sHMA) domain containing proteins as potential targets of some of these effectors, including Pwl2 (Zdrzałek *et al*., 2024). Small HMAs are highly expanded across plant species with putative functions including heavy metal detoxification and potential metallochaperones, but in most cases their functions are not known (Dykema *et al*., 1999; Suzuki *et al*., 2002; Gao *et al*., 2009). HMAs can be broadly classified into two families, heavy metal-associated plant proteins (HPPs), and heavy metal-associated isoprenylated plant proteins (HIPPs), which possess a C-terminal isoprenylation motif (CaaX, where ‘a’ represents an aliphatic residue and ‘X’ is any amino acid) important for membrane anchoring (Hála and Žárský, 2019). The rice sHMA protein Pi21, for example, is a blast disease susceptibility factor (Fukuoka *et al*., 2009), which has led to deployment of loss-of-function alleles of *pi21*, as a recessive form of rice blast resistance (Fukuoka *et al*., 2009; Mutiga *et al*., 2021). Furthermore, differences in host sHMA protein repertoires are linked to host-specific, adaptive evolution of the APikL2 effector family (Bentham *et al*., 2021). Importantly, HMA protein domains have been identified as non-canonical immune sensory domains in some NLR proteins, such as the rice Pik-1 of the Pik-1/2 and RGA5 of the RGA5/4 pair receptors respectively, essential for recognition of cognate *M*. *oryzae* effectors (Ashikawa *et al*., 2008; Kanzaki *et al*., 2012; Cesari *et al*., 2013; Maqbool *et al*., 2015). However, the function of sHMAs and their link to disease susceptibility is not understood, limiting our understanding of the role of MAX effectors in plant disease.

In this study we set out to investigate the biological function of *PWL2*. We were motivated to understand how a broadly distributed effector such as Pwl2 functions during a susceptible interaction between *M. oryzae* and its host. We report that *PWL2* has undergone extensive duplication and expansion in copy number with many rice blast isolates possessing 3-5 copies, making its functional analysis challenging. Using CRISPR-Cas9 gene editing, however, we have generated a *Δpwl2* mutant confirming that Pwl2 is a host specificity determinant but also revealing a hitherto un-recognized virulence function. Furthermore, we demonstrate here that *PWL2* expression during infection is controlled by the Pmk1 MAP kinase, which regulates cell-to-cell movement by the fungus at plasmodesmata-containing pit fields (Sakulkoo et al., 2018). We reveal that Pwl2 targets the HIPP43 sHMA protein in barley, thereby displacing it from plasmodesmata and show that transgenic plants overexpressing either *PWL2* or HIPP43 have reduced immune responses and greater susceptibility to blast disease. Finally, we show that a Pwl2^SNDEYWY^ mutant unable to interact with HIPP43 fails to displace it from PDs and cannot complement the reduced virulence of *Δpwl2* mutants. When considered together, our study provides evidence that Pwl2 is a virulence factor in *M. oryzae* that suppresses host defense by re-localising HIPP43 away from PDs to facilitate fungal invasion of plant tissue.

## Results

### Pwl2 is a cytoplasmic effector expressed during blast infection

Expression of *PWL2* is specific to the initial biotrophic phase of plant infection, peaking at 48 hours post-infection (hpi) (Yan *et al*., 2023) (**Fig. S1A - C**). The Pwl2 effector localises in the biotrophic interfacial complex (BIC), a plant membrane-rich structure –that is clearly visible as a single bright punctum, initially at the tip of a penetration hypha and then adjacent to bulbous, branched invasive hyphae (Khang *et al*., 2010). The BIC is the predicted site of effector delivery (Kankanala *et al*., 2007; Giraldo *et al*., 2013; Oliveira-Garcia *et al*., 2023b; Were and Talbot, 2023), although experiments to date have not precluded that the BIC could be a site of effector sequestration from the host plant. To investigate this possibility, we generated a single *M. oryzae* strain expressing two BIC-localised effectors Pwl2-mRFP and Bas1-GFP and visualised their localisation during infection of leaf sheath tissue in the susceptible rice cultivar Moukoto. The two effectors were observed as small punctate signals within the same BIC (**Fig. 1A and B**). By contrast, when two different *M. oryzae* strains, Ina168 and Guy11, expressing Pwl2-GFP and Pwl2-mRFP respectively were used to simultaneously infect rice tissue, we observed that the BIC formed by each individual invasive hypha exclusively contained either Pwl2-GFP or Pwl2-mRFP respectively but never both fluorescence signals (**Fig. 1C and D)**. This is consistent with Pwl2 secretion by each fungal strain into the BIC, because sequestration of previously secreted Pwl2 from the plant cytoplasm to the BIC would result in a mixed GFP/mRFP fluorescence signal in the BIC. We conclude that Pwl2 is expressed early during infection and secreted to the BIC from where it is delivered into host cells.

**Fig. 1.**
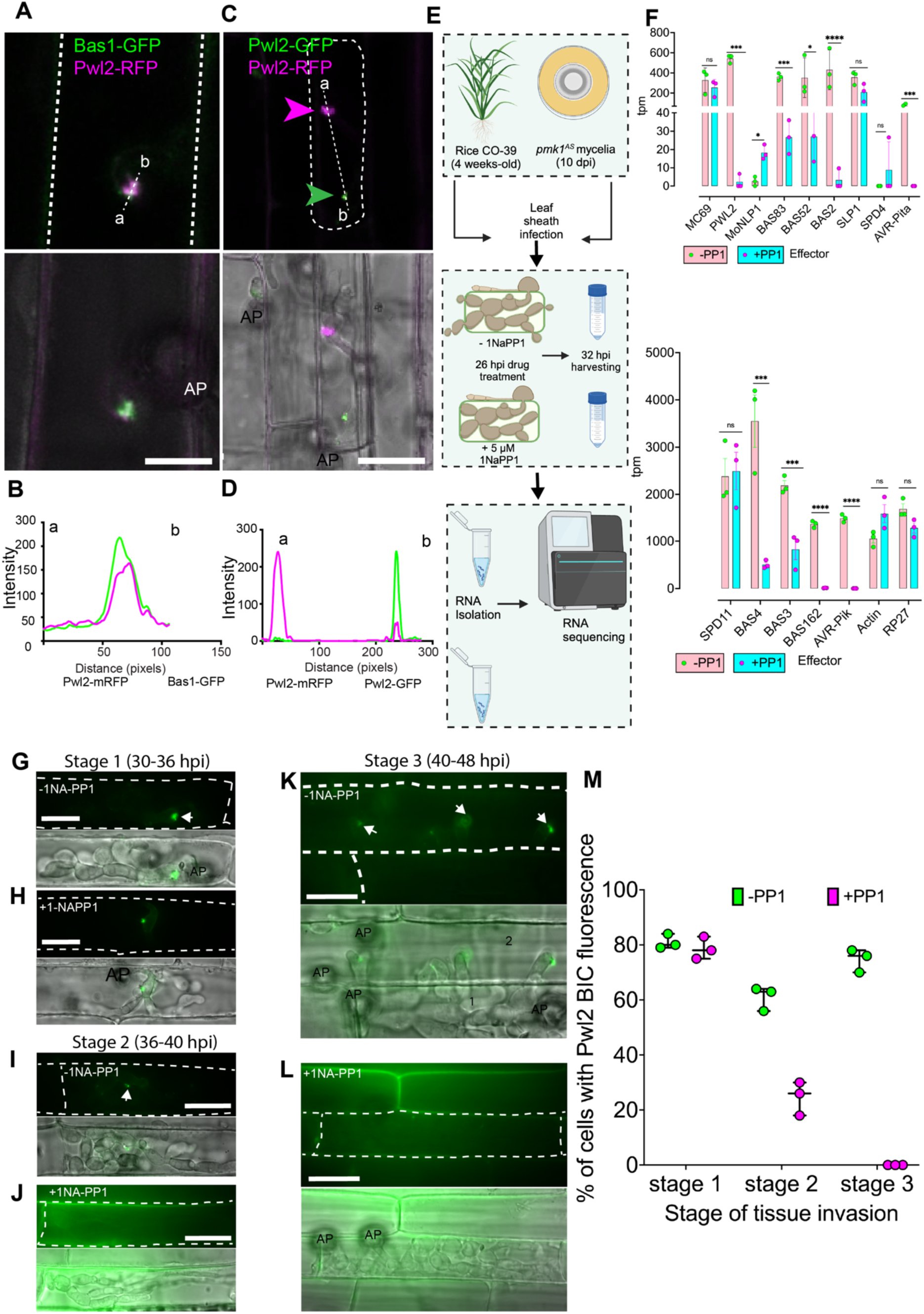
*PWL2* expression is regulated in a Pmk1-dependent manner during host infection. (**A+B**) Micrographs and line scan graph showing Pwl2 secreted through the Biotrophic Interfacial Complex (BIC) in rice cells during early infection. Conidial suspension at 1x 10^5^ mL^-1^ of *M. oryzae* strain co-expressing two BIC localised effectors, Bas1-GFP and Pwl2-mRFP were inoculated onto a susceptible cultivar Moukoto rice leaf sheath and images captured at 26 hpi. Fluorescence of the two effectors was observed as small punctate signals in the same BIC. (**C+D**) Co-infection assay of rice leaf sheath with two different *M. oryzae* strains, one expressing Pwl2-mRFP and the other Pwl2-GFP at 30 hours post-infection (hpi). Micrographs and line scan graph shows there is an absence of mixed fluorescence confirming the BIC does not contain Pwl2 transferred from rice cells. BICs indicated by magenta arrows for mRFP and green arrows for GFP. (**E**) Schematic illustration to describe the workflow used to test genes regulated in a Pmk1-dependent manner. Rice leaf sheaths were infected with a *M. oryzae pmk1^AS^* mutant spores before mock or 1NA-PP1 treatment. Treated and mock treated, infected leaf sheaths were trimmed and used for RNA isolation followed by sequencing. Figure created with BioRender https://biorender.com/. (**F**) Bar charts to show that a subset of effectors is regulated in a manner that requires Pmk1. Gene expression is shown as TPM (transcripts per million) at 32 hpi. (**G-L**) Micrographs showing expression of Pwl2-GFP by *M. oryzae pmk1^AS^* in leaf sheaths of a susceptible rice line CO39 using conidial suspension at 1x 10^5^ mL^-1^. Fluorescence of Pwl2 at different stages of infectious hyphal progression, starting with early stage of infection, Stage 1 (30-36 hpi) where a newly differentiated bulbous hyphae is formed as per (Sakulkoo *et al*., 2018). Analysis was also carried out at later stages of infection including Stage 2 (36-40 hpi) where a primary invaded cell is filled with differentiated bulbous hyphae) and Stage 3 (40-48 hpi) where colonization of primary invaded cell is complete and there is full invasion of secondary invaded cells. To test Pmk1 inhibition, inoculated leaf sheaths were treated with 5 µM 1NA-PP1 at 26 hpi. (**M**) Inhibition of Pwl2 by 1NA-PP1 is quantified as percentage of cells showing Pwl2 fluorescence in the BIC. Arrows indicate fluorescence in the BIC. Scale bars represent 20 µm. The primary and secondary invaded cells are indicated with 1 and 2 respectively. Significance between groups of samples was performed using Unpaired Student’s t-test. \*\*\*\**P*<0.0001, \*\**P*<0.01, \**P*<0.05, NS indicates no significant difference.

### *PWL2* expression is regulated by the Pmk1 MAP kinase during invasive growth

The blast fungus invades rice tissue by means of pit field sites containing plasmodesmata (PDs), which enable it to move between rice cells while maintaining integrity of the rice plasma membrane. PD conductance is also regulated by the fungus, enabling effectors like Pwl2 to be deployed in adjacent uninfected cells (Kankanala et al., 2007; Sakulkoo et al., 2018). Cell-to-cell movement by the fungus is regulated by the pathogenicity mitogen-activated protein kinase 1 (Pmk1 MAPK) pathway. An analogue-sensitive mutant of Pmk1 has been shown to be unable to move through pit fields in the presence of the MAPK inhibitor 1NA-PP1 (Sakulkoo et al., 2018). Given that *PWL2* is expressed during the initial stages of infection, we decided to test whether it is regulated by Pmk1. We therefore re-analyzed RNA-seq data (**Fig. 1E**) (Sakulkoo *et al*., 2018) by separating raw reads of *M. oryzae* and *O. sativa* and quantifying transcript abundance using Kallisto, followed by determining differential expression using Sleuth. We found that *PWL2* is significantly down-regulated in a *M. oryze pmk1^AS^*mutant during cell-to-cell movement in the presence of 1NA-PP1 together with a subset of known effector genes - *BAS83*, *BAS52*, *BAS2, AVR-Pita-1, BAS3, BAS4, BAS162* and *AVR-Pik-C* (**Fig. 1F**). By contrast, *MoNLP1* showed up-regulation, while expression of actin (MGG_03982) and the RP27 40S 27a ribosomal subunit genes (MGG_02872) were not significantly affected (**Fig. 1F**). We then used live cell imaging to investigate Pwl2-GFP expression in a *pmk1^AS^*mutant during rice leaf sheath infection ±1Na-PP1. We initially observed Pwl2-GFP in the BIC at early stages of infection, unaffected (Stage 1) 30-36hpi (**Fig. 1G and H**), as previously reported (Sakulkoo *et al*., 2018), but by later stages of infection Stage 2 (36-40hpi), (**Fig. 1I-J**), and Stage 3 (40-48hpi) (**Fig. 1K-L**), 14-22h after 1NA-PP1 treatment, Pwl2-GFP fluorescence was significantly reduced (**Fig. 1J, L and M**). The reduction in Pwl2-GFP fluorescence was consistently associated with the stage at which *M. oryzae* traverses PD-containing pit field sites and enters adjacent plant cells. Taken together, we conclude that Pwl2 is regulated by the Pmk1 MAPK signalling pathway during cell-to-cell invasive growth by *M. oryzae*.

### *PWL2* is highly conserved in *M. oryzae*

*PWL2* belongs to a large gene family (Kang *et al*., 1995; Sweigard *et al*., 1995), but its conservation in the global rice blast population is not known. We therefore investigated *PWL* gene family distribution in isolates that infect a variety of different grass species (**Fig. 2A**) This revealed that *PWL2* is found in the majority of host-limited forms of *M. oryzae* and related *Magnaporthe* species, except for *Setaria* and closely-related *Panicum, Cynodon* and *Urochloa*-infecting isolates (although these were less well represented than other isolates) as shown in **Fig. 2A**. By contrast, *PWL1* is present in a sub-set (Asuke *et al*., 2020) of *Eleusine*-infecting isolates and some *Oryza*-infecting isolates, but largely absent from other host-specific lineages, except one *Eragrostis*-infecting isolate (EtK19-1), and one *Cynodon*-infecting isolate (Cd88215) (**Fig. 2A)**. Similarly, *PWL3* is present in most *Oryza*-infecting isolates, some *Setaria*, *Lolium*, and *Eleusine*-infecting isolates, but largely absent in *Eragrostis, Triticum*, *Digitaria* and *Pennisetum* lineages, while *PWL4* is present in *Eleusine*, *Eragrostis, Lolium* and *Triticum*-infecting isolates, *Pennisetum* and *Digitaria*-infecting isolates, but only two *Oryza*-infecting isolates (**Fig. 2A)**. We conclude that the *PWL* gene family (Kang *et al*., 1995) is broadly distributed among blast fungus isolates infecting numerous grasses but that *PWL2*, is widespread across rice-infecting isolates and the majority of host-limited forms of *M. oryzae*.

**Fig. 2.**
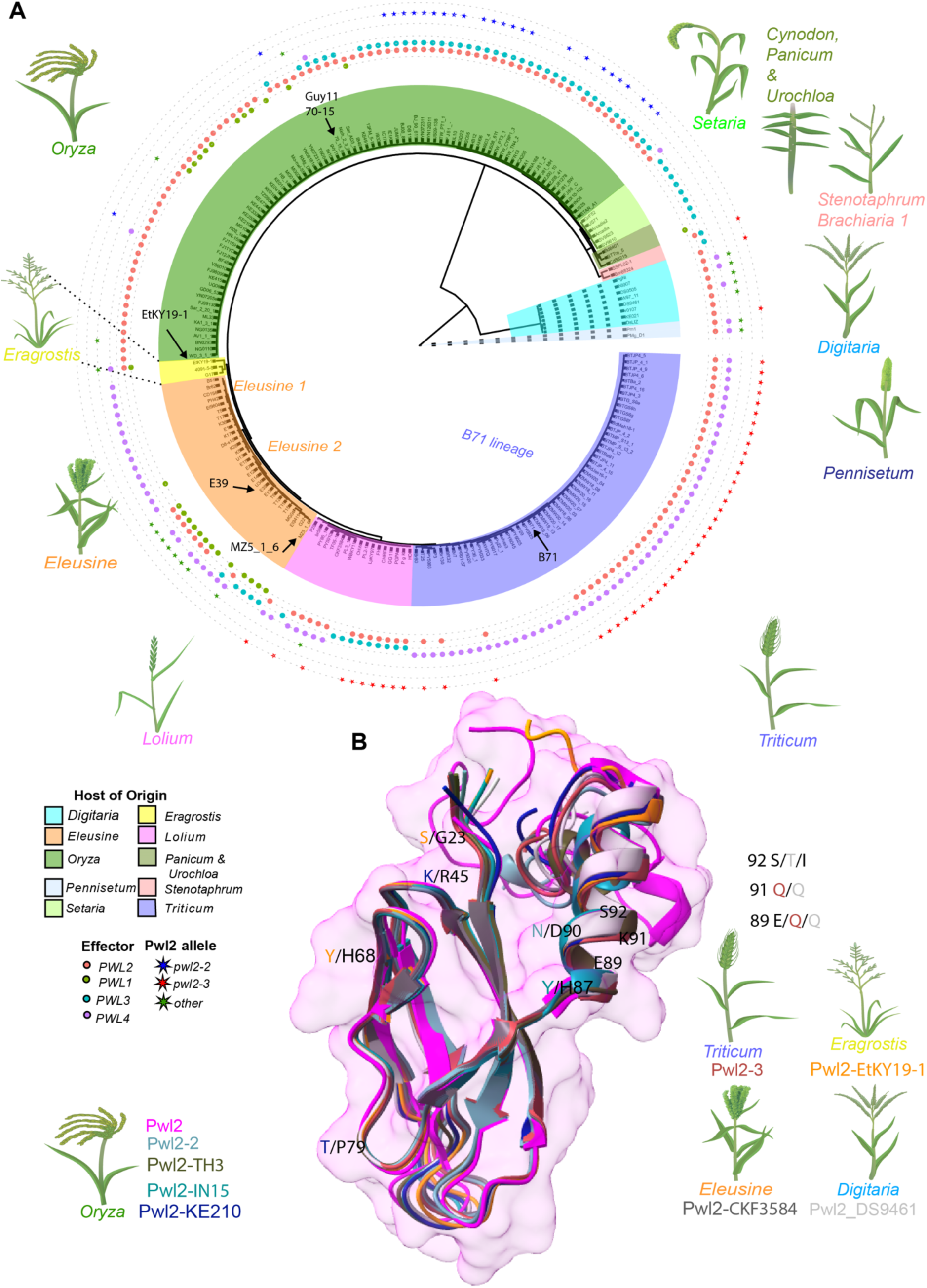
Pwl2 is highly conserved in isolates of *M. oryzae* and displays conserved structural features. (**A)** Phylogenetic analysis and *PWL* gene family distribution in *M. oryzae*. A maximum parsimony tree was generated using kSNP3 to include isolates from different host-limited forms of *M. oryzae* including isolates that infect *Oryza sativa* (rice), *Eleusine* spp. (finger millet), *Hordeum vulgare* (barley), *Setaria* spp. (foxtail millet), *Triticum aestivum* (wheat), *Lolium* spp. (rye grass), *Brachiaria* spp. (signal grass), *Panicum* spp. (torpedo grass), *Eragrostis* spp. (weeping lovegrass), *Stenotaphrum* spp. (buffalo grass), *Cynodon* spp. (Bermuda grass) and *Urochloa* spp. (signal grass) (Ou, 1980; Talbot, 2003; Cruz and Valent, 2017; Inoue *et al*., 2017), as well as *Magnaporthe* species that infect *Digitaria sanguinalis* (crabgrass) and *Pennisetum* spp. (pearl millet). We used Pwl1 (BAH22184.1), Pwl2 (QNS36448.1), Pwl3 (AAA80240.1) and Pwl4 (AAA80241.1) protein sequences to query the presence or absence of each gene using tblastn. The heatmap indicates the presence/absence of genes in the *PWL* family. *PWL1* is predominantly present in group EC-1I (Asuke *et al*., 2020) of *Eleusine*-infecting isolates and some *Oryza*-infecting isolates, but largely absent from other host-specific lineages, except one *Eragrostis*-infecting isolate, EtK19-1, and one *Cynodon*-infecting isolate, Cd88215, but not in *Digitaria* and *Pennisetum* spp lineages. *PWL3* is present in most *Oryza*-infecting isolates, some *Setaria*, *Lolium*, and *Eleusine*-infecting isolates but largely absent in *Eragrostis* and *Triticum* lineages, as well as *Digitaria* and *Pennisetum* lineages. *PWL4* is present in *Eleusine*, *Eragrostis, Lolium* and *Triticum*-infecting isolates, but only two *Oryza*-infecting isolates. Several *Pennisetum* and *Digitaria*-infecting isolates, however, carry *PWL4. PWL2* is found in most host-limited forms of *M. oryzae* and related *Magnaporthe* species but absent in *Brachiaria, Setaria, Panicum, Cynodon* and *Urochloa*. (**B**) Superimposition of different Pwl2 variants predicted using AlphaFold3 onto the resolved Pwl2 structure (magenta) indicating region of polymorphism. Different color represent different variant as follows, Pwl2-2 (teal), Pwl2-3 (brick red), Pwl2-TH3 (dark olive green), Pwl2-IN15 (dark teal), Pwl2-KE210 (blue), Pwl2-CKF3584 (dark grey), Pwl2-EtKY19-1 (orange), Pwl2-DS9461 (light grey). The superimposition shows overall structural conservation in MAX-fold, without the signal peptide and the C-terminus. The variants of Pwl2 are named with corresponding isolate name and grouped according to host species. Varying residues are colored according to the genome from which they were identified.

Having determined that *PWL2* is broadly distributed in *M. oryzae*, we next sought to determine its allelic variability. In addition to a loss of recognition allele *pwl2-2*, which contains a single Asp-90-Asn substitution (Sweigard *et al*., 1995), we identified 14 new alleles of *PWL2* (**Fig. S2A**). Notably, most polymorphic residues occur between positions His87 to Ser92, suggesting these residues might contribute to Pwl2 recognition by a cognate resistance gene (**Fig. S2A**). Interestingly, we could only identify 1 variant of *PWL1*, 5 variants of *PWL3* and 1 variant of *PWL4*, despite these effector genes occurring in finger millet, rice, wheat and ryegrass-infecting lineages, respectively (**Fig. 2A and S2B - S2D**). It is possible that sample size for some host-specific lineages may lead to an underestimate of allelic variability in *PWL1, PWL3*, and *PWL4,* but from this analysis *PWL2* appears to be highly polymorphic and conserved, compared to other members of the gene family (**Fig. S2E**). We also tested the ability of a subset of *PWL2* alleles to be recognised by barley *Mla3*. In some cases, we found variant *PWL2* alleles occurred in a *M. oryzae* isolate carrying *PWL1* (such as U34 and E39) or isolates with multiple copies of *PWL2* (for example TH3), precluding functional analysis, and not all isolates were available for testing. As expected, Guy11 did not produce lesions on cv. Baronesse, but was able to infect cv. Nigrate (**Fig. S3A – C**). However, JUM1 (*pwl2-2*), BTJP4-16 (*pwl2-3)* and BN0293 (variant *pwl2*) all caused blast disease on cv. Baronesse, confirming that they contain loss of recognition *pwl2* alleles (**Fig. S3A - C**). Pwl2 has recently been structurally described as a MAX effector, containing a core β-sandwich fold formed of two antiparallel β-sheets, as well as a single α-helix, C-terminal to the β-sandwich fold (Zdrzałek *et al*., 2024). We were interested to understand how these variable residues affect Pwl2 recognition. We therefore mapped a subset of the identified variants of Pwl2 to the effector crystal structure. We noted that all polymorphic residues in *pwl2* alleles, such as the Glu-89-Gln, Lys-91-Gln and Ser-92-Ile substitutions in *pwl2-3,* were present in the C-terminal α-helix, a distinct region from the core MAX-fold (**Fig. 2B**). We conclude that this interface is likely used by immune receptors present in resistant barley (Mla3) and weeping lovegrass to bind Pwl2 and is potentially stabilized by the MAX-fold.

### *PWL2* has undergone copy number expansion in *M. oryzae*

*PWL2* copy number appears to vary significantly among *M. oryzae* isolates. The *M. oryzae* reference genome assembly (strain 70-15) has two copies of *PWL2* on chromosomes 3 and 6, annotated as MGG_04301 and MGG_13683, respectively (Dean *et al*., 2005). Southern blot hybridization (**Fig. S4**), however, and analysis of long-read assembled genome sequence of *M. oryzae* Guy11 identified three copies of *PWL2* (**Fig. 3A).** *PWL2* loci were associated with *MGR583*, *POT2* and *MAGGY* transposable elements, suggesting their potential involvement in genome rearrangements (**Fig. 3A)**. To further assess *PWL2* copy number variation in *M. oryzae* isolates, we employed k-mer analysis to determine copy number variation of *PWL2* in 286 *M. oryzae* genomes. Copy number variation is common among effector-encoding genes, such as *AVR-Pik*, *AVR-Pizt, BAS1, BAS4* and *SLP1*, but was particularly prevalent for *PWL2* with one isolate, for instance, containing 9 copies (**Fig. 3B).** We conclude that *PWL2* has expanded in copy number in *M. oryzae*.

**Fig. 3.**
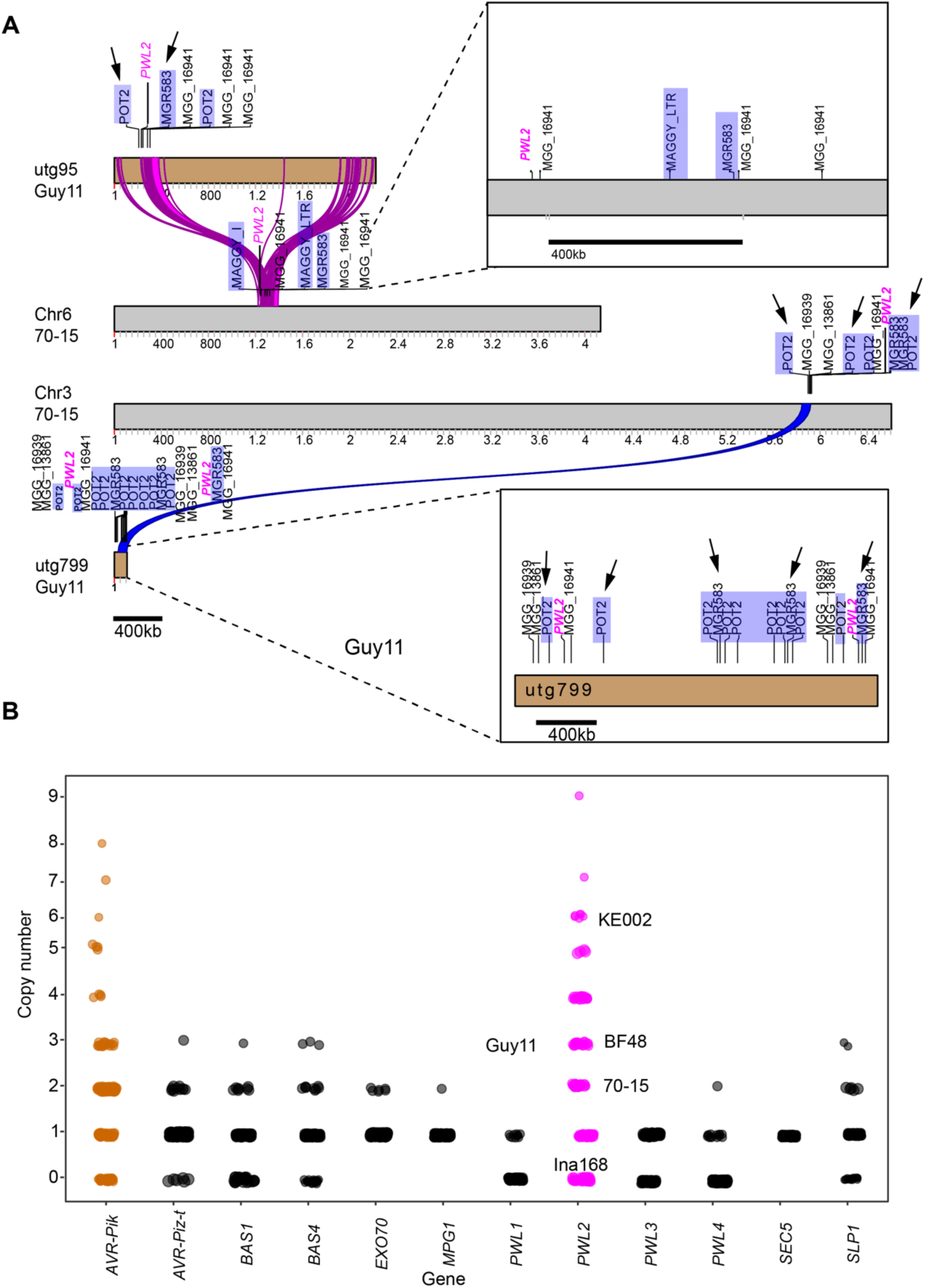
*PWL2* has undergone copy number expansion in *M. oryzae* field isolates. **(A)** Schematic diagram showing the estimated chromosomal location of *PWL2* on multiple loci (Chr3 and 6) of the reference genome 70-15 and on equivalent region of assembled contigs of laboratory strain Guy11 genome. *PWL2* is flanked by *POT2* and *MGR583* repeated sequences suggesting a possible involvement in translocation of events into different loci in the genome. Arrows indicate location of *POT2* and *MGR58* while *PWL2* is labelled in magenta. (**B)** A k-mer analysis on sequenced raw reads was used to determine copy number variation of *PWL* family genes in different *M. oryzae* isolates. Plot shows high copy number of *PWL2* in analyzed genomes (n = 286) compared to the other *PWL* gene family members, another effector *SLP1,* and selected control genes *MPG1* and *SEC5*. Similarly, *AVR-Pik* shows multiple copies in different isolates. Copy number of *PWL2* in selected isolates Guy11, KE002, BF48, 70-15 is indicated with Ina168 that lack *PWL2* used as a negative control.

### Pwl2 is both a host range determinant and a virulence factor

We next set out to determine the function of *PWL2* in blast disease through gene functional analysis. Given that *PWL2* occurs in multiple copies, we used CRISPR/Cas9 genome editing (Foster *et al*., 2018) to simultaneously delete all three copies of *PWL2* found in Guy11. We designed a sgRNA to target *PWL2* and introduced a hygromycin phosphotransferase (*HPH*) encoding gene cassette (**Fig. 4A)**. Transformants with ectopic integrations, or where some *PWL2* had not been deleted, gave a predicted PCR amplicon of 645 bp in size. By contrast, *Δpwl2* mutants generated a larger amplicon of 1.5 kb (**Fig. 4A**). Four putative transformants were selected for whole genome sequencing. Raw reads were aligned to the *M. oryzae* reference genome using samtools v.15, before converting into BAM files and visualising using IGV viewer. No reads mapping to the *PWL2* locus were identified in mutants T6 and T12, whereas fewer reads mapped to the same locus from sequenced T4 and T5 (**Fig. S5A**). This confirmed that T6 and T12 are *Δpwl2* transformants. Transformant T4, T5 and T6 displayed vegetative growth that was identical to Guy11 with normal dark concentric rings and light growing edges, while T12 showed slight reduced growth, melanisation and conidiation (**Fig. S5B)**. Given that *PWL2* is a host range determinant and avirulence gene, we reasoned that *M. oryzae Δpwl2* deletion mutants would gain virulence on weeping lovegrass (Kang *et al*., 1995; Sweigard *et al*., 1995) and barley cv. Baronesse expressing *Mla3* (Brabham *et al*., 2023). The *Δpwl2* deletion transformants T5, T6 and T12 and Guy11 were therefore used to infect weeping lovegrass seedlings and barley Baronesse using spray inoculation. The *Δpwl2* mutant T6 produced disease symptoms identical to those produced by *Eragrostis*-infecting isolate G17 and *Oryza*-infecting isolate Ina168 which lacks *PWL2,* while Guy11 and the two complemented isogenic strains (T6+*PWL2*p:*PWL2*) and (T6+*RP27*p:*PWL2*) were unable to cause disease (**Fig. 4B**). Similarly, T5 and T12 also produced disease symptoms on weeping lovegrass seedlings and cv. Baronesse (**Fig. S5C**). We conclude that T5, T6 and T12 are loss of recognition mutants of *PWL2* (**Fig. 4B and S5C**), and we can be confident that T6 and T12 have complete deletion of all three copies of the gene.

**Fig. 4.**
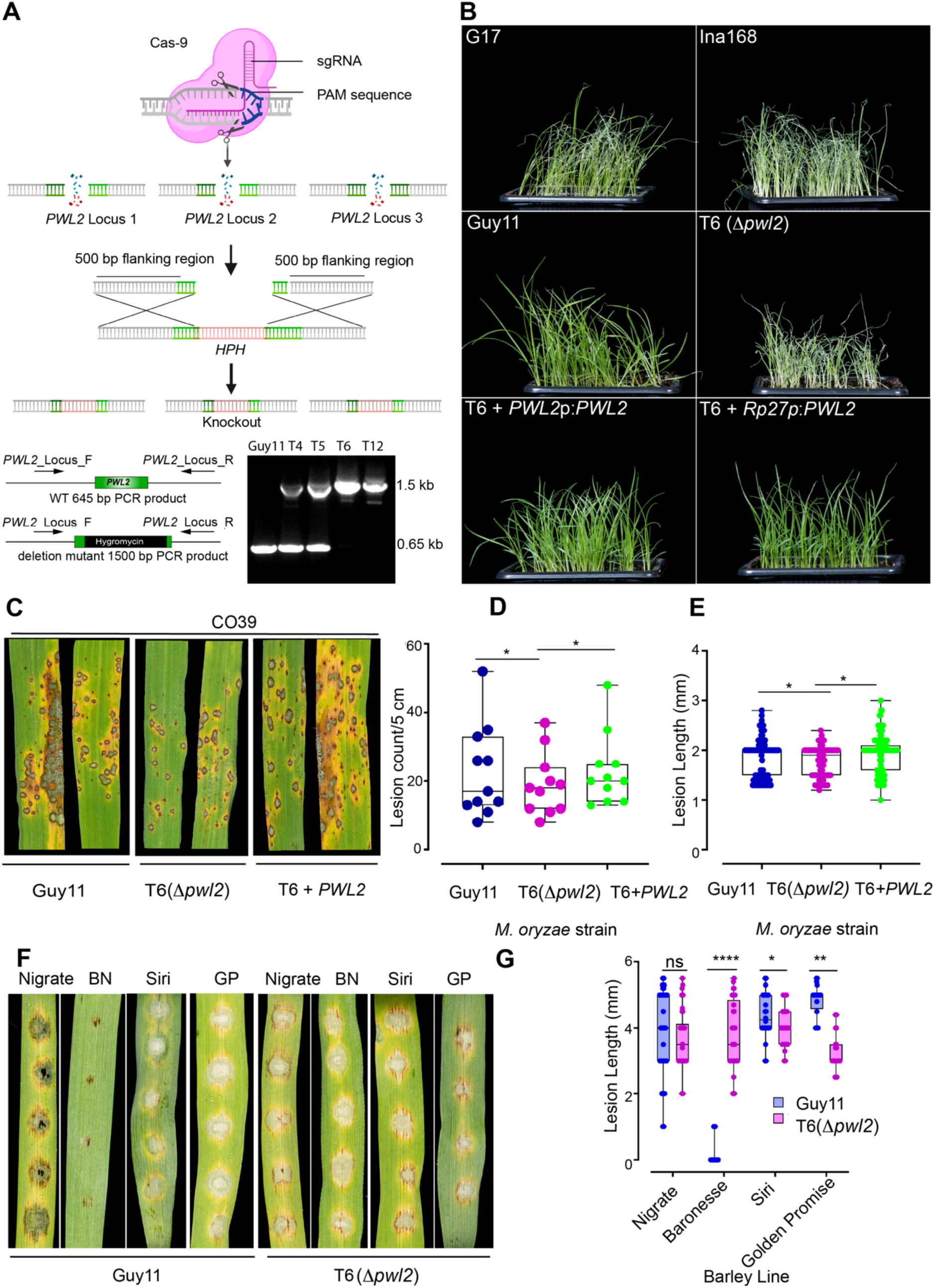
CRISPR-Cas9 *Δpwl2* mutants demonstrate Pwl2 as both host range and a virulence factor **(A)** Schematic illustration of the CRISPR-Cas9 mediated gene editing process for inserting the Hygromycin Phosphotransferase (*HPH*) gene cassette at the *PWL2* locus. The guided sequence (sgPWL2) directs Cas9 to introduce a double stranded break at the *PWL2* locus, with the DNA repair template containing the Hygromycin resistance gene cassette and flanking regions of the *PWL2* gene. This leads to mutations in the form of indels or gene replacements. Figure created with BioRender https://biorender.com/. Positive mutants were identified by amplifying the hygromycin cassette from transformants with the three copies deleted. (**B)** Comparison of disease symptoms in weeping lovegrass (*E. curvula*) infected with different *M. oryzae* isolates. In the top panel, typical disease lesions produced by *E. curvula*-infecting isolate G17 (*-PWL2*) and a rice infecting isolate Ina168 (*-PWL2*) are shown. In the middle panel, CRISPR-Cas9 deletion mutants exhibit gain of virulence on weeping lovegrass compared to Guy11. In the bottom panel, complemented strains with both native and constitutive RP27 promotors showed a loss of virulence on weeping lovegrass. (**C**) *Δpwl2* mutants display reduced pathogenicity on rice cultivar CO39. Conidial suspensions from Guy11, *Δpwl2* (T6) and complemented *Δpwl2*(T6) +*PWL2*p:*PWL2*) were used to inoculate 21-day-old seedlings of the blast-susceptible cultivar CO39, and disease symptoms recorded after 5 dpi. The box plots show the lesion density (**D**) and lesion size (**E**) of seedlings infected with Guy11, *Δpwl2* (T6) and complemented *Δpwl2* (T6 +*PWL2*p:*PWL2*). (**F**) The *Δpwl2* mutant exhibits enhanced pathogenicity on barley cultivar Baronesse (*Mla3*). Conidial suspensions from Guy11 and *Δpwl2* T6 were used to inoculate 10-day old seedlings of barley lines Golden Promise, Siri, Nigrate and Baronesse, and disease symptoms recorded after 5 dpi. Conidial suspensions at 1 ×10^5^ mL^−1^ spores/mL was used for infection assays. **(G)** The box plot shows lesion size of barley seedlings infected with Guy11 and *Δpwl2* (T6). The lower horizontal line shows the minimum value, and the upper horizontal line shows the maximum value. The lower border and upper border of the box shows the lower quartile and upper quartile, respectively. The line in the box shows the median. Significance between groups of samples was performed using Unpaired Student’s t-test. \*\*\*\**P*<0.0001, \*\**P*<0.01, \**P*<0.05, NS indicates no significant difference.

We next tested whether Pwl2 has a virulence function during a compatible interaction between the blast fungus and its host. Having observed that T12 had slight differences in vegetative growth, we decided to select T6 for virulence assays and as a genetic complementation background for *PWL2*. T6 was used to inoculate susceptible rice cultivar CO39 by spray infection, and on susceptible barley lines cv. Nigrate (-*Mla3)*, Siri (+*Mla8*), cv. Golden Promise (+*Mla8*), and resistant cv. Baronesse (+*Mla3*) using leaf drop inoculation. In four independent replicates, the *Δpwl2* mutant T6 showed a reproducible reduction in virulence on CO39 compared to wildtype Guy11 or a complemented isogenic strain (T6+*PWL2*p:*PWL2*), based on lesion density and lesion length (**Fig. 4C - E)**. As expected, T6 was able to cause blast disease on cv. Baronesse unlike Guy11 and produced statistically smaller lesion sizes on barley lines Siri and Golden Promise, compared to Guy11. Both strains were able to infect Nigrate (**Fig. 4F and G**). We conclude that *PWL2* contributes to the ability of *M. oryzae* to cause blast disease.

### Pwl2 suppresses host immunity

Having determined that the Pwl2 effector contributes to virulence, we decided to study its biological function during host cell colonisation. To do this we first generated stable transgenic barley lines (in cv. Golden Promise) expressing *PWL2-YFP* (without its signal peptide) under control of the CaMV35S promoter (**Fig. 5A)**. We tested two independent *PWL2-YFP* transgenic plants for their response to two elicitors of PAMP-triggered immunity (PTI), flg22 and chitin, compared to wild type cv. Golden Promise (**Fig. 5A)**. Barley perceives flg22 through the pattern recognition receptor FLS2 and chitin through HvCEBiP and HvCERK1, leading to immune responses such as generation of reactive oxygen species (ROS) (Yu *et al*., 2023). We observed that Pwl2 expression abolished flg22 (**Fig. 5B - C)** and chitin-induced ROS generation (**Fig. 5D - E)**. Pwl2 therefore contributes to virulence by suppressing PTI during compatible interactions. To investigate the effect of elevated Pwl2 effector expression on blast infection, we infected two independent transgenic barley plants expressing *PWL2* with *M. oryzae* Guy11. *PWL2*-expressing barley lines were more susceptible and developed blast disease symptoms earlier (2-3 dpi) compared to infection of isogenic cv. Golden Promise (**Fig. 5F and G)**. We conclude that Pwl2 acts as a modulator of PTI that helps facilitate fungal infection.

**Fig. 5.**
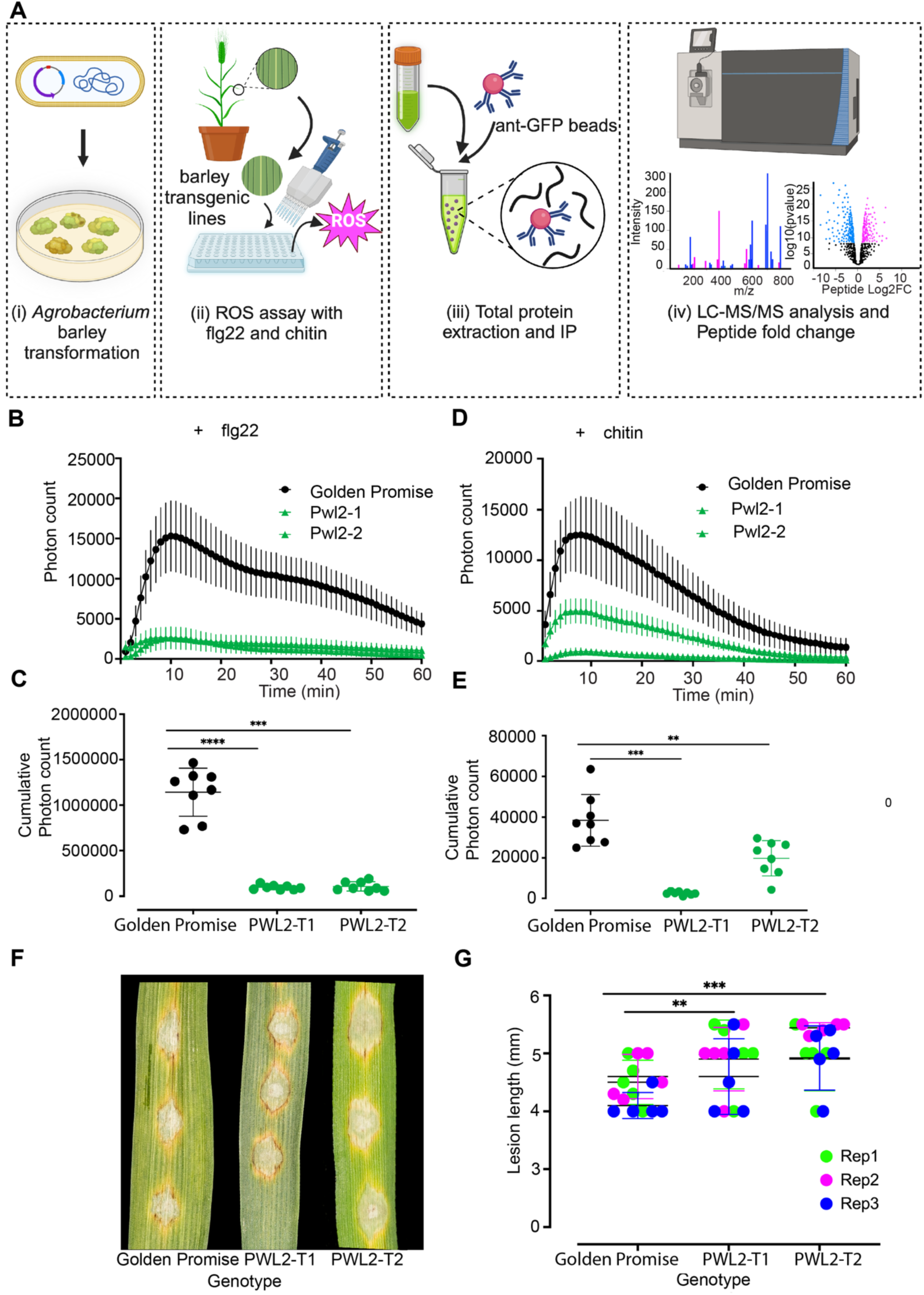
Pwl2 suppresses PAMP-induced ROS in transgenic barley lines (**A**) Schematic illustration to describe the workflow used to generate transgenic plants, test for PTI response and identify putative Pwl2 interactors using discovery proteomics. ROS production in leaf discs collected from 4–week-old stable transgenic lines expressing Pwl2-YFP compared to cv. Golden Promise induced by 1 uM flg22 (**B-C**) or chitin (**D-E**) (n > 8). Points represent mean; error bars represent SEM. For ROS assay, line graphs (**B+D**) points represent mean per time point and dot plots represent cumulative ROS (**C+E**) production over 60 min, error bars represent SEM. For dot plot, graphs show the average cumulative photon count for cv. Golden Promise control and two independent Pwl2-YFP lines. The lower horizontal line shows the minimum value, and the upper horizontal line shows the maximum value, the middle line shows the mean value. (**F)** Leaf drop infection on two independent barley transgenic lines expressing Pwl2-YFP compared to infection on wild type cv. Golden Promise 3-4 dpi. (**G**) The dot plot represents the disease lesion length on cv. Golden Promise compared to two independent Pwl2-YFP transgenic lines. Unpaired Student’s t-test was performed to determine significant differences \*\*\*\**P*<0.0001, \*\*\**P*<0.001, \*\**P*<0.01. These experiments were repeated three times to obtain consistent result.

### Pwl2 interacts with the heavy metal binding isoprenylated protein HIPP43

To determine the likely target of Pwl2, transgenic barley lines expressing Pwl2-YFP were used in co-immunoprecipitation coupled to mass spectrometry (IP-MS) analysis. This was followed by spectral search in the *H. vulgare* (barley, cv Morex version 3) proteome database. We identified a total of 282 frequently occurring proteins in the eight biological replicates containing Pwl2, that were not identified in the six free-YFP control samples. To avoid focusing on false positives in the form of sticky and abundant proteins, fold-changes were calculated by first determining the average number of peptides per protein candidate, before estimating log_2_ fold change compared to the control experiment. We further filtered for proteins that produced at least two peptide hits in more than half of replicates (≥4/8 biological replicates). A total of 52 proteins met this criterion and were considered as enriched in samples compared to controls and were therefore selected for further analysis. Furthermore, because Pwl2 attenuates the ROS-burst in barley transgenic lines, we focused initially on protein candidates previously reported to have potential roles in immunity or ROS generation (**Fig. 6A)**. These were selected for one-to-one interactions using a yeast-two-hybrid (Y2H) assay by co-transforming constructs expressing Pwl2 and <u>P</u>utative <u>P</u>wl2 <u>I</u>nteracting <u>P</u>roteins (PPIPs) identified by IP-MS (**Fig. 6A)**. Pwl2 showed a strong interaction with PPIP4, a HMA-domain protein (**Fig. 6B)**. We named this protein *Hv*HIPP43, for *Hordeum vulgare* heavy metal domain containing isoprenylated plant protein 43 based on homology to *Os*HIPP43 (Zdrzałek *et al*., 2024). We also tested to ensure that neither Pwl2 nor HIPP43 demonstrated auto-activity in yeast-2-hybrid drop out media (**Fig. S6A**). To independently verify the interaction between Pwl2 and HIPP43, we carried out co-immunoprecipitation (co-IP) analysis using protein extracts from *N. benthamiana* and confirmed that *Hv*HIPP43 interacts with Pwl2-YFP (**Fig. 6C and D)**. Conversely, three other MAX-fold effectors, MEP3, AVR-PikE, AVR-Piz-t, or free YFP did not interact with HIPP43 (**Fig. 6D)**. We next tested whether *PWL* gene family-encoded proteins Pwl1, Pwl3, Pwl4 and the pwl2-3 variant could also interact with *Hv*HIPP43 and showed that they can all interact based on Y2H assays, suggesting that Pwl2 belongs to an effector family that interacts with a host sHMA protein (**Fig. 6E)**. By contrast, three MAX-fold effectors AVR-Pia, AVR-PikD and AVR-Mgk did not interact with HIPP43 (**Fig. 6E)**. Barley and wheat have three copies of HIPP43 per haploid genome, but based on our IP-MS results, we identified peptides that mapped to two copies of HIPP43, which both strongly interacted with Pwl2 in Y2H assays HIPP43 (**Fig. S6B-C)**.

**Fig. 6.**
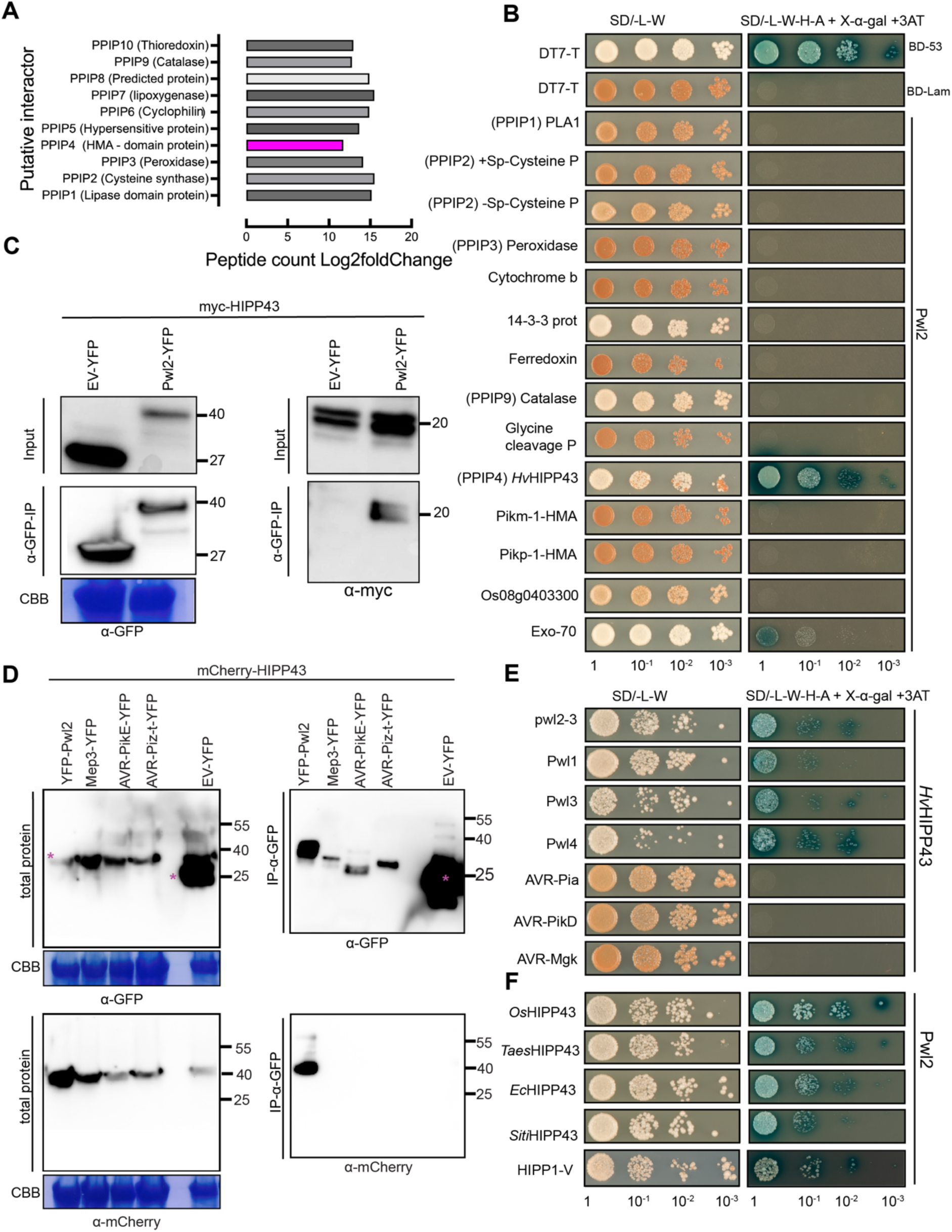
Pwl2 interacts with HIPP43 and its orthologs from other grass species. (**A**) Putative Pwl2 interacting peptides were immunoprecipitated from protein extracts of 3-week-old stable cv. Golden Promise transgenic lines expressing Pwl2-YFP or free cytoplasmic YFP using anti-GFP antibodies, and LC-MS/MS was performed to identify unique putatively interacting peptides. The scale 0-20 represents the Log2fold change in peptides when compared to the control. (**B**) One-to-one yeast two-hybrid between Pwl2 and selected top candidates from IP-MS analysis, Pwl2 and <u>P</u>utative <u>P</u>wl2 Interacting <u>P</u>roteins (PPIPs). HMA integrated in rice NLR Pikm-1 and Pikp-1 were used as specificity controls. Simultaneous co-transformation of pGADT7-Pwl2 (prey vector) and pGBK-PPIPs (bait vector) and PGBKT7-53 and pGADT7-T (positive control) pGADTT7-T and pGBKT7-Lam (negative control) into Y2H gold strain. Positive interaction resulted in the activation of four reporter genes and growth on high-stringency medium (−Ade, −Leu, –Trp, _His +X-α-gal and 3-amino-1,2,4-triazole). Co-transformation also activates the expression of *MEL1*, which results in the secretion of α– galactosidase and the hydrolysis of X-α-gal in the medium, turning the yeast colonies blue. HIPP43 exclusively interacts with Pwl2 in SD/-L-W-H-A X-α-gal medium with 3AT added. These experiments were repeated several times over three years obtaining consistent result. **(C**) Co-immunoprecipitation (co-IP) of Pwl2-YFP and Myc-HIPP43 or (**D**) mCherry-HIPP43 in *N. benthamiana* leaves. C-or N-terminal GFP tagged Pwl2 and C-terminal Myc-HIPP43 was cloned into the vector pGW514 and transformed into *Agrobacterium* strain *GV3101* and co-infiltrated into *N. benthamiana* leaves and left to incubate for 48 h. Immunoprecipitates were obtained with anti-GFP affinity matrix beads and probed with anti-GFP-peroxidase, anti-mCherry-peroxidase and anti-Myc-peroxidase (HRP-conjugated) antibodies. Total protein extracts were also probed with appropriate (HRP-conjugated) antibodies. (**E**) Yeast-two-hybrid analysis shows; upper panel, *Hv*HIPP43 interacts with *PWL* gene family products Pwl1, Pwl3, Pwl4 and variant *pwl2-3*. (**F**) Pwl2 interacts with HIPP43 homologs from rice (*Os*HIPP43), wheat (*Taes*HIPP43), weeping love grass (*Ec*HIPP43), foxtail millet (*SitiHIPP43*) and wild wheat (HIPP1-V). These experiments were repeated several times over three years obtaining consistent result.

### Pwl2 can interact with HIPP43 orthologs of diverse grass species

HIPPs are expanded in plant genomes and to test whether HIPP43 has orthologs in other grass species, we generated a maximum likelihood phylogenetic tree of HMA domains from diverse grasses with high quality genomes and annotations (**Fig. S6D**). The HIPP43 family formed a distinct clade which includes orthologs from rice (*O. sativa)*, wheat (*T. aestivum*), wild wheat (*Haynaldia villosa*), foxtail millet (*S. italica*), weeping lovegrass (*E. curvula*), Sorghum (*Sorghum bicolor*), *Oropetium thomaeum*, *Zea mays* and *Brachypodium distachyon* (**Fig. S6D**). Furthermore, we could identify copy number variation ranging from one to five copies in grasses, with barley and wheat (i.e. *Triticeae* lineage) having three or more paralogs of HIPP43, one per haploid genome (**Fig. S7A**). We found that Pwl2 interacts with *Hv*HIPP43 orthologs from rice (*Os*HIPP43), *Triticum* (*Taes*HIPP43), *Setaria* (*SitiHIPP43*), *Eragrostis* (*Ec*HIPP43) and *Haynaldia* (HIPP1-V) in Y2H assays (**Fig. 6F)**. The interaction of Pwl2 with rice *Os*HIPP43 was verified by *in vitro* biochemistry and a structure of the complex was obtained by *X*-ray crystallography (Zdrzałek *et al*., 2024). We also analyzed the transcriptional profile of *Os*HIPP43 during infection of Guy11 in a susceptible rice line CO39 (Yan *et al*., 2023) and found that *Os*HIPP43 expression is up-regulated during plant infection, consistent with a role in host defense (**Fig. S7B)**.

### Overexpression of HIPP43 suppresses PTI and increases blast susceptibility

To investigate the role of the Pwl2-HIPP43 interaction, we generated barley transgenic lines overexpressing *Hv*HIPP43. Two independent transgenic lines were challenged with two PTI elicitors, flg22 (**Fig. 7A-B)** and chitin (**Fig. 7C-D)**. Strikingly, we observed that both the chitin and flg22-induced ROS burst was abolished in plants expressing *Hv*HIPP43, compared to wild type (**Fig. 7A-D)**. Similarly, when we challenged transgenic barley plants expressing *Hv*HIPP43 with *M. oryzae*, we observed increased susceptibility compared to the wildtype cv. Golden Promise, and disease lesions appeared earlier than in wild type plants (**Fig. 7E and F)**. These findings are consistent with over-expression of both *PWL2* and HIPP43 leading to enhanced blast disease susceptibility.

**Fig. 7.**
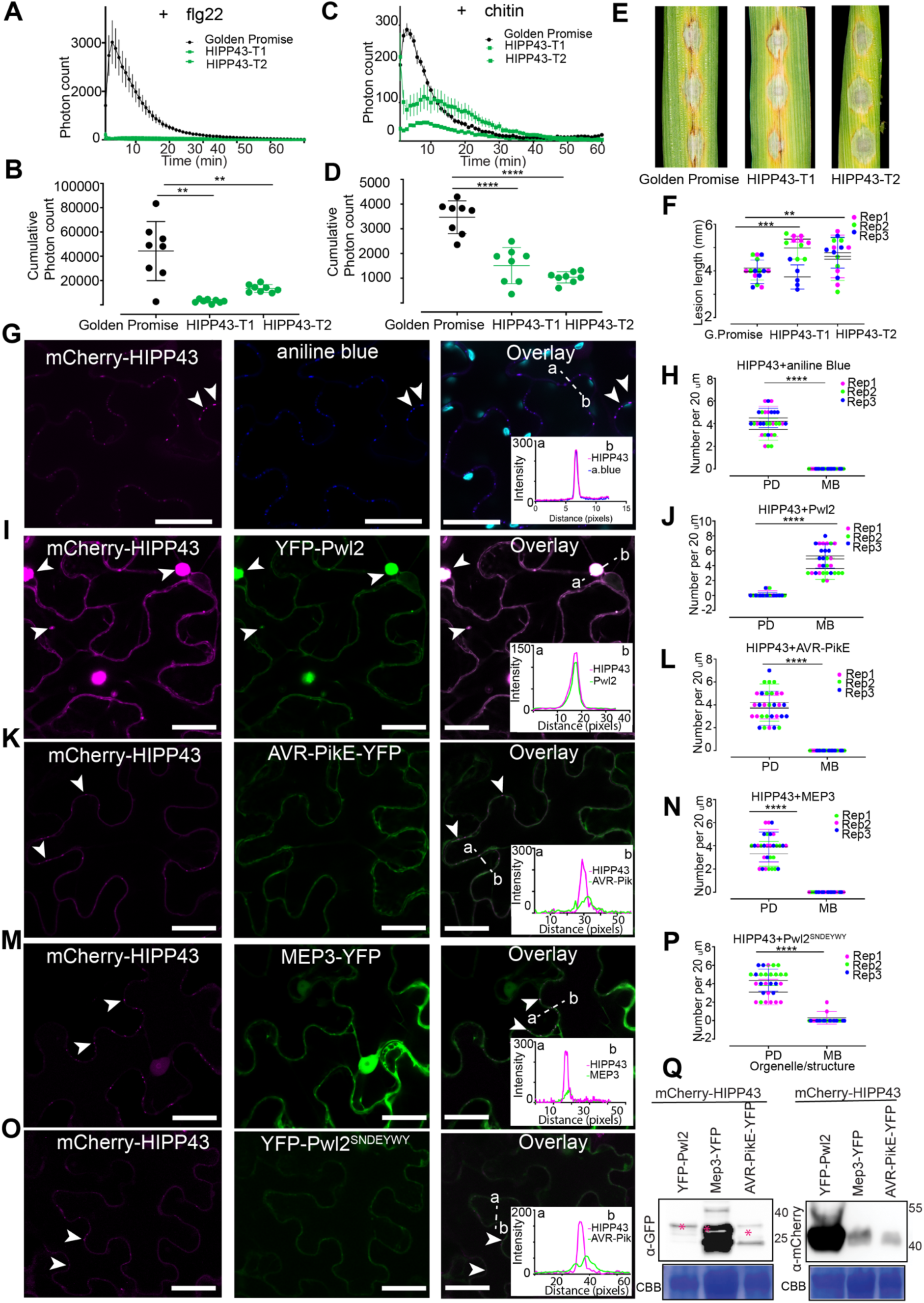
*Hv*HIPP43 suppresses PAMP-induced ROS in transgenic barley and is stabilized by Pwl2. **(A-D)** ROS production measured from leaf disks collected from 4–week-old stable transgenic line expressing YFP-HIPP43 and cv. Golden promise (control) in the absence and presence of 1 uM flg22 or 1 mg/ml Chitin (n > 8). For ROS assay, line graphs (**A+C**) points represent mean per time point and dot plots represent cumulative ROS (**B+D**) production over 60 min, error bars represent SEM for cv. Golden Promise control and two independent YFP-HIPP43 lines. For dot plot, graphs show the average cumulative photon count for cv. Golden Promise control and two independent YFP-HIPP43 lines. (**E**) Leaf drop infection on barley transgenic lines expressing YFP-HIPP43 compared to wild type Golden Promise. Conidial suspensions at 1 ×10^5^ mL^−1^ spores/mL from Guy11 were used for inoculation. Disease symptoms were recorded after 4 dpi. (**F**) Dot plot showing lesion length on barley lines infected with Guy11. All experiments were repeated three times giving consistent results. **(G+H)** Micrographs and line scan graph showing mCherry-HIPP43 localizing as small puncta on the plasma membrane when expressed in *N. benthamiana.* Staining of callose using aniline blue overlaps with mCherry-HIPP43, confirming mCherry-HIPP43 localises exclusively at the plasmodesmata (PD) localization and cytoplasmic mobile bodies (MB) localization is absent. **(I+J)** Micrographs and line scan graph showing the presence of YFP-Pwl2, alters mCherry-HIPP43 localisation, which was observed to translocate to the cytoplasm, mobile cytoplasmic bodies. **(K+L)** Micrographs and line scan graph showing mCherry-HIPP43 remains in the PD in the presence of cytoplasmically localised AVR-PikE**. (M+N**) Micrographs and line scan graph showing mCherry-HIPP43 remains in the PD in the presence of cytoplasmically and PD localised MEP3. (**O+P**) Micrographs and line scan graph showing mCherry-HIPP43 remains in the PD in the presence of Pwl2^SNDEYWY^, a mutant that does not interact with HIPP43. Dotted lines in the overlay panels correspond to distance and direction of line intensity plot. Scale bars represent 20 µm. For dot plots, the lower horizontal line shows the minimum value, the upper horizontal line shows the maximum value, the middle line shows the mean value. Unpaired Student’s t-test was performed to determine significant differences \*\*\*\**P*<0.0001, \*\*\**P*<0.001, \*\**P*<0.01. (**Q**) Western blot to show that Pwl2 does not degrade HIPP43. Total protein extracts from *N. benthamiana* leaves co-infiltrated with YFP-Pwl2, MEP3 or AVR-PikE and mCherry-HIPP43 were immunoblotted and probed with anti-GFP-peroxidase (left) or anti-mCherry-peroxidase (right) (HRP-conjugated) antibodies. Microscope imaging experiments were repeated several times over two years giving consistent result.

### Pwl2 prevents plasmodesmatal localisation of HIPP43

To investigate the function of *Hv*HIPP43, we performed live cell imaging following transient expression in *N. benthamiana* to determine its sub-cellular localisation. When tagged at the N-terminus (mCherry-HIPP43), *Hv*HIPP43 localised predominantly to puncta at the plasma membrane that co-localise with callose staining (aniline blue), consistent with sites of plasmodesmata (PD) (**Fig. 7G and H)**. Like all HIPPs, *Hv*HIPP43 has an isoprenylation motif at its C-terminus (**Figure S8A and B**), implicated in membrane anchoring (Hála and Žárský, 2019), so we investigated whether PD localisation is affected by its deletion. We observed that PD localisation was indeed abolished in the absence of the isoprenylation motif and-Iso-mCherry-HIPP43 (*Hv*HIPP43 without isoprenylation motif) instead accumulated in the nucleoplasm and at the plasma membrane. Moreover, co-expression with a plasma membrane marker LTi6b-GFP and aniline blue staining confirmed its localisation at the plasma membrane and loss of PD localisation (**Fig. S8C)** compared to the wild type (**Fig. S8D)**.

Given the strong interaction between Pwl2 and HIPP43 and the localisation of HIPP43 to PD, we were keen to see the effect of co-expressing the effector and putative host target in the same cells. When we co-expressed YFP-Pwl2 with mCherry-HIPP43, they co-localised in the cytosol as mobile puncta approximately 2 μm in diameter (**Fig. 7I**). In some cases, we observed larger mobile structures ranging from 2-4 μm in diameter (**Fig. 7I**). We also tested whether Pwl1, Pwl3 and Pwl4 (**Fig. S9A**) could also co-localise with HIPP43 when co-expressed in *N. benthamiana.* In the presence of Pwl1, HIPP43 PD localization was abolished, and the two proteins co-localise in the cytoplasm and partially in cytoplasmic mobile structures (**Fig. S9B**) Conversely, Pwl3 did not affect mCherry-HIPP43 localisation to PD (**Fig. S9C**), while Pwl4 (**Fig. S9D**) only showed cytoplasmic co-localisation without larger mobile bodies forming. In addition, because we had observed that *Hv*HIPP43 orthologs from wheat (*Taes*HIPP43), foxtail millet (*Siti*HIPP43) and weeping lovegrass (*Ec*HIPP43) interact with Pwl2 in Y2H assays, we also tested whether *Taes*HIPP43, *Siti*HIPP43 or *Ec*HIPP43 localise as PD puncta in the same way as *Hv*HIPP43. Interestingly, *Taes*HIPP43 (wheat) mostly localized to the nucleus, cytoplasm and PD, even though PD localization was reduced (**Fig. S9E**), whereas *Ec*HIPP43 (weeping lovegrass) (**Fig. S9F**) and *Siti*HIPP43 (foxtail millet) **(Fig. S9G**) localized to small puncta equivalent to PD, like *Hv*-HIPP43. Because Pwl2 is able to alter sub-cellular localisation of *Hv*HIPP43, we tested whether this also occurred when *Siti*HIPP43 and Pwl2 were co-expressed. We observed Siti-HIPP43/Pwl2 co-localisation as cytoplasmic mobile bodies away from PD, mirroring the Pwl2/HvHIPP43 interaction (**Fig. S9H-J**). When considered together, we conclude that Pwl2 and its family members can interact with *Hv*HIPP43 and that Pwl2 can also interact with HIPP43 orthologs from other cereals, thereby altering their deployment to PD.

### Pwl2 consistently alters the plasmodesmatal localization of HIPP43

To understand the effect of Pwl2 expression on the PD localisation of *Hv*HIPP43, we carried out a detailed quantitative analysis. In control experiments we expressed *Hv*HIPP43 with free cytoplasmic YFP, and two MAX effectors AVR-PikE and MEP3 (Yan *et al*., 2023). In six independent co-expression experiments, we observed co-localisation of Pwl2 and *Hv*HIPP43 as mobile cytoplasmic puncta or large cytoplasmic mobile structures, while PD localisation was significantly reduced (**Fig. 7I and J)**. By contrast, co-localisation of *Hv*HIPP43 with AVR-PikE or MEP3 led to mCherry-*Hv*HIPP43 fluorescence remaining at PDs (**Fig. 7K*-*N)**. Moreover, mCherry-HIPP43 fluorescence showed greater intensity when co-expressed with Pwl2 compared to when co-expressed with AVR-PikE or MEP3 **(Fig. 7K-N)**. It is therefore possible that Pwl2 is able to stabilise *Hv*HIPP43 or increases its accumulation, because the mCherry-*Hv*HIPP43 signal is barely detectable in the absence of Pwl2 (**Fig. 7K and M)**. Our previous structural analysis demonstrated that Pwl2 and *Os*HIPP43 produce a robust binding interface that requires up to seven mutations in Pwl2 (*Pwl2*^SNDEYWY^) to abolish (Zdrzałek *et al*., 2024). To test whether binding of Pwl2 to HIPP43 is necessary for the alteration in its cellular localisation, we co-infiltrated mCherry-HIPP43 and YFP-Pwl2^SNDEYWY^ in *N. benthamiana* and carried out live cell imaging. We were able to observe YFP-Pwl2^SNDEYWY^ expression, but this did not alter mCherry-*Hv*HIPP43 localisation at PD (**Fig. 7O-P)**. Furthermore, the fluorescence intensity of mCherry-*Hv*HIPP43 did not increase as it did upon co-expression with YFP-Pwl2 **(Fig. 7I and O)**. To rule out the possibility that Pwl2 degrades *Hv*HIPP43 upon interaction in plant cells, we carried out immuno-blot analysis using protein extracts from *N. benthamiana* following co-infiltration of mCherry-HIPP43 with either Pwl2, MEP3 or AVR-PikE. Strikingly, the presence of Pwl2 led to pronounced mCherry-*Hv*HIPP43 accumulation compared to when co-infiltrated with MEP3 or AVR-PikE **(Fig. 7Q)**. We were also concerned that Pwl2/HIPP43 co-localisation as puncta could be mis-interpreted as also occurring at PD. To rule out such a possibility, we stained PD with aniline blue after co-infiltrating *Hv*HIPP43 with free-YFP, YFP-Pwl2, AVR-PikE-YFP and MEP3-YFP. In a control experiment we expressed free-YFP, YFP-Pwl2, AVR-PikE-YFP and MEP3-YFP in the absence of *Hv*HIPP43. We found that free-YFP, Pwl2, AVR-PikE-YFP and MEP3 show localisation to the cytosol **(Fig. S10A-D**), but MEP3-YFP also showed some additional PD localisation. By contrast YFP-Pwl2 was always observed as cytoplasmic mobile puncta (**Fig. S10B**). When cells expressing Pwl2 were stained with aniline blue there was no co-localization of Pwl2 and the aniline blue signal (**Fig. S10E-G**). In the presence of mCherry-HIPP43 and YFP-Pwl2, co-localisation at cytoplasmic puncta/mobile bodies was observed and did not co-localise with aniline blue staining (**Fig. S10H**). In limited cases where co-localisation was observed, this was transient because the Pwl2/HIPP43 puncta are mobile. By contrast, when expressed on its own, or in the presence of free-YFP or two MAX-fold effectors AVR-PikE, free-YFP and MEP3, mCherry-HIPP43 co-localised with aniline blue to PD (**Fig. S10I-K**). To investigate the motility of HIPP43-Pwl2 puncta, we carried out time-lapse imaging using confocal laser microscopy of aniline blue-stained *N. benthamiana* cells following co-infiltration of mCherry-HIPP43 with YFP-Pwl2, MEP3-YFP, AVR-PikE-YFP or YFP-Pwl2^SNDEYWY^. Blue stained puncta equivalent to PD remained immobile while mCherry-HIPP43/YFP-Pwl2 fluorescence was observed at mobile cytoplasmic structures (see **Movie S1** and individual frames shown in **Fig. S11A)**. By contrast, co-localisation of mCherry-HIPP43 at aniline blue-stained PD was clearly visible and not altered by the presence two MAX-fold effectors MEP3-YFP and AVR-PikE-YFP, YFP-Pwl2^SNDEYWY^ or of free-YFP (see **Movie S2-S5**) and frames in **Fig S11B-E)**. When considered together, these results provide evidence that Pwl2 interacts with HIPP43 altering its cellular location away from plasmodesmata.

### The ability of Pwl2 to bind HIPP43 is necessary for its avirulence and virulence functions

As the interaction between Pwl2 and HIPP43 appears to be critical for altering its sub-cellular localisation to PD, we decided to test if this was also necessary for the role of Pwl2 in blast disease. We therefore set out to see whether the *pwl2*^SNDEYWY^ allele could complement the mutant phenotypes of a *M. oryzae Δpwl2* mutant. To do this, we transformed *Δpwl2* mutant with a construct expressing Pwl2^SNDEYWY^ under its native promoter (*PWL2p:pwl2^SNDEYWY^*) and successful transformants were selected (**Fig. 8A and B**). We then quantified *Pwl2^SNDEYWY^* gene expression in selected transformants using qRT-PCR (**Fig. 8C**). We reasoned that if recognition of Pwl2 by Mla3 requires HIPP43 recognition (Gómez De La Cruz *et al*., 2024), then Pwl2^SNDEYWY^ should not complement the observed gain of virulence of *Δpwl2* mutants on barley cv. Baronesse (+*Mla3*). Moreover, if the Pwl2/HIPP43 interaction is required for the virulence function of Pwl2, then Pwl2^SNDEYWY^ should not restore virulence to *Δpwl2* mutants. In three independent experiments of two independent strains, C6 (*Δpwl2*+*PWL2p:pwl2^SNDEYWY^*) and C14 (*Δpwl2*+*PWL2p:pwl2^SNDEYWY^*) showed a compatible interaction on cv. Baronesse (**Fig. 8D)** providing evidence that the Pwl2^SNDEYWY^ is not recognised by Mla3, because the strains retained the ability to infect a susceptible barley cv. Nigrate (-*Mla3*) (**Fig. 8E)**. We also observed that *Δpwl2* +*PWL2p:Pwl2^SNDEYWY^* transformants C6 and C14 did not restore full virulence on susceptible rice cv. CO39 compared to the wildtype Guy11 (**Fig. 8F**), based on lesion density (**Fig. 8G)** and lesion length (**Fig. 8H)**. We conclude that the interaction between Pwl2 and HIPP43 is required for both its recognition as an avirulence factor and its function as a virulence determinant.

**Fig. 8.**
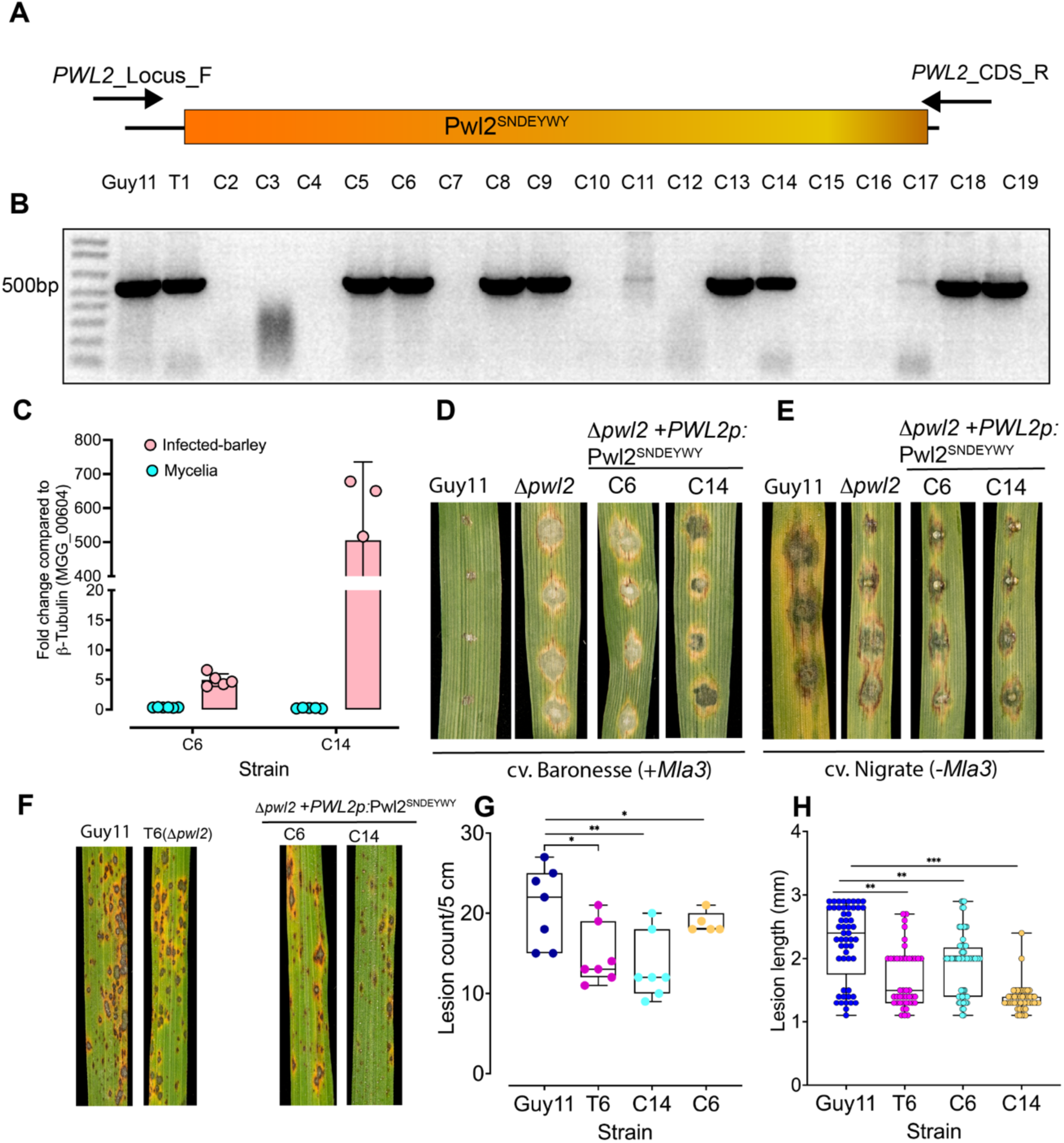
Pwl2^SNDEYWY^ does not complement Mla3 recognition and virulence on a blast-susceptible rice cultivar CO39. (**A and B**) Complemented *Δpwl2*+*PWL2*p:*Pwl2^SNDEYWY^*) positive transformants were screened using PCR to amplify *PWL2* coding sequence. (**C**) Bar charts showing relative expression as log2 fold change of *Pwl2^SNDEYWY^* in two selected transformants, C6 and C14 using qRT-PCR. Detached leaves of 10-day old seedlings of barley were inoculated with *Δpwl2*+*PWL2*p:*Pwl2^SNDEYWY^*) and infected tissue collected 40 hpi and used for RNA isolation (3 biological replicates), cDNA synthesised and samples used for qRT-PCR. (**D and E**) *Δpwl2*+*PWL2*p:*Pwl2^SNDEYWY^*complemented strains C6 and C14 produced compatible disease lesions on barley cultivar Baronesse (+*Mla3*) (**D**) and Nigrate (-*Mla3*) (**E**). Conidial suspensions at 1 ×10^5^ mL^−1^ spores/mL from Guy11, *Δpwl2* and complemented *Δpwl2*+*PWL2*p:*Pwl2^SNDEYWY^* were used to inoculate 10-day old seedlings of barley and disease symptoms recorded after 5 dpi. (**F-H**) Complemented *Δpwl2*+*PWL2*p:*Pwl2^SNDEYWY^*display reduced pathogenicity on rice cultivar CO39. Conidial suspensions at 1 ×10^5^ mL^−1^ spores/mL from Guy11, *Δpwl2* T6 and complemented *Δpwl2*+*PWL2*p:*Pwl2^SNDEYWY^* were used to inoculate 21-day-old seedlings of the blast-susceptible cultivar CO39, and disease symptoms recorded after 5 dpi. Significance between groups of samples was performed using Unpaired Student’s t-test. \*\*\**P*<0.0001, \*\**P*<0.01, \**P*<0.05, NS indicates no significant difference.

## Discussion

The Pwl2 effector was first identified as a host specificity determinant for infection of the forage grass species weeping lovegrass (Kang *et al*., 1995; Sweigard *et al*., 1995). Its initial identification provided evidence that host range in plant pathogenic fungi was conditioned in a similar way to cultivar specificity in a gene-for-gene manner, involving dominant pathogen genes recognised by the products of cognate disease resistance genes (Kang *et al*., 1995). Furthermore, Pwl2 was found to belong to an expanded gene family, suggesting an important function in pathogenicity and fitness (Kang *et al*., 1995). However, in the following two decades, the function of Pwl2 remained elusive despite its extensive use as a marker in cell biological studies for investigating effector regulation, secretion and delivery during plant infection (Kankanala *et al*., 2007; Giraldo *et al*., 2013; Oliveira-Garcia *et al*., 2023b).

In this study, we set out to explore the function of Pwl2 and investigate why it is such a highly conserved effector in *M. oryzae*. We found that *PWL2* is highly conserved in *M. oryzae*, including each of its host-limited forms and even in sister *Magnaporthe* species infecting crabgrass and pearl millet. The observation that *PWL2* and the wider *PWL* family genes have been maintained in the global blast population and there are effector variants in certain host-adapted lineages, suggests that Pwl2 and members of this family serve an important function in pathogenesis. Furthermore, *PWL2* is present at high copy number in many *M. oryzae* isolates having undergone extensive gene duplication and is specifically expressed during the initial stages of blast infection, particularly at the stage when *M. oryzae* moves from an initially colonised epidermal cell, following appressorium penetration, to adjacent host cells. This cell-to-cell movement by the fungus utilises PD-containing pit fields and the fungus forms a transpressorium structure that undergoes severe hyphal constriction to traverse each pit field (Kankanala *et al*., 2007; Cruz-Mireles *et al*., 2021), a process that requires activity of the Pmk1 MAP kinase (Sakulkoo et al., 2018). We found that *PWL2* is expressed during this process in a Pmk1-dependent manner, forming part of a regulated set of effectors deployed by the fungus during cell-to-cell movement. The generation of a *Δpwl2* mutant, made possible by CRISPR/Cas9 gene editing to delete all three native copies of the gene, enabled us to confirm the role of Pwl2 in host specificity to weeping lovegrass (Sweigard et al., 1995), as an avirulence effector for barley Mla3 (Brabham et al., 2024), and also revealed its importance in blast disease. Pwl2 is therefore an important virulence determinant for blast disease, explaining its conservation and amplification.

To identify the likely target of Pwl2, we utilised discovery proteomics which revealed its interaction with an isoprenylated small HMA protein, HIPP43. This is consistent with Pwl2 being a MAX effector – many of which have been shown to interact with sHMA protein domains – although previous studies have focused on incorporation of sHMA domains into paired NLR immune receptors leading to disease resistance (Ortiz *et al*., 2017; Bentham *et al*., 2021; Maidment *et al*., 2021; Mukhi *et al*., 2021). The crystal structure of the Pwl2/HIPP43 complex (Zdrzałek *et al*., 2024) demonstrates that Pwl2 uses an expansive interface to mediate binding to HIPP43, largely using elements of the MAX fold to interact with the β-sheet of its host target. Indeed, when HIPP43 is incorporated into the Pik-1 NLR, in place of its naturally occurring integrated HMA domain, this leads to an immune response to Pwl2 (Zdrzałek *et al*., 2024). A recent study has also revealed that the barley resistance protein Mla3 acquired the ability to bind Pwl2 by mimicking the HMA fold of its host target, HIPP43 (Gómez De La Cruz *et al*., 2024). Interestingly, polymorphic residues found in Pwl2 variants, such as pwl2-2 and pwl2-3 are not located at the HIPP43 binding interface but are located away from the MAX fold in the C-terminal helix of Pwl2. It is possible that these polymorphic residues are essential to stabilize the Pwl2, MAX-fold/HMA-like interface in Mla3. Consistent with this idea, introducing mutations that disrupt the Pwl2/HIPP43 interaction also results in loss of recognition by Mla3.

In spite of the importance of sHMA domains in plant immunity, little is known regarding their actual function or why they are targeted by fungal effectors. HMA domains are known to be involved in biotic/abiotic stress responses, transport of metals and metal detoxification, and their expression can be organ-specific or tissue-specific within roots, leaves, or stems (Barth *et al*., 2009; Zschiesche *et al*., 2015; Zhang *et al*., 2020). Moreover, sHMAs can localize to the nucleus, plasma membrane, cytoplasm, or to plasmodesmata (Barth *et al*., 2009; Cowan *et al*., 2018; Barr *et al*., 2023; Oikawa *et al*., 2024). Furthermore, sHMAs are expanded in plant genomes with more than 45, and more than 50, sHMA domain-encoding genes occurring in *Arabidopsis* and rice, respectively (de Abreu-Neto *et al*., 2013). Unique intragenic deletions in OsHIPP05 (*Pi21*), a proline rich HMA domain protein-encoding gene in rice leads to rice blast resistance (Fukuoka *et al*., 2009) and gene silencing or deletion mutants of *TaHIPP1* or *AtHMAD1* in wheat and *Arabidopsis* provide enhanced resistance against *Puccinia striiformis* f. sp*. tritici* and *Pseudomonas syringae* DC3000, respectively (Imran *et al*., 2016; Wang *et al*., 2023). In addition, deletion mutants of *AtHIPP27* in *Arabidopsis* lead to increased resistance against cyst nematodes (Radakovic *et al*., 2018). However, it is not clear why deletion of sHMA protein-encoding genes impacts immunity. Therefore, the observation that Pwl2 interacts with HIPP43 is revealing, especially because over-expression of either Pwl2 or *Hv*HIPP43 suppresses PTI responses and enhances blast disease susceptibility. How this potential enhancement of HIPP43 activity is stimulated by Pwl2 is not completely clear, but transient co-expression of both Pwl2 and HIPP43 sequesters HIPP43 away from plasmodesmata (PD), and the two proteins instead co-localize in large mobile structures within the cytoplasm. The localisation of HIPP43 to PD requires its isoprenylation motif and may be associated with an immune signaling role at this site. Given the Pmk1 MAP kinase-dependent expression of Pwl2 during PD traversal, it may be that sequestering HIPP43 away from these sites is critical for the fungus to invade plant tissue efficiently. This is consistent with the reduced virulence phenotype of *Δpwl2* mutants, which results in slower generation of disease symptoms and reduced lesion size. A recent study has highlighted how a *P. infestans* effector PiE354 can interfere with a host immune response by re-routing a plant Rab8a away from the plant/pathogen interface, a similar potential effector function that alters the cellular location of a target rather than impairing a particular function (Yuen *et al*., 2024).

Localization of two HIPP proteins, HIPP7 and HIPP26, to PD has been previously reported, which is also dependent on isoprenylation motifs like HIPP43 (Cowan *et al*., 2018; Guo *et al*., 2021). Although characterized HIPPs have been proposed to be metallochaperones, there are no studies that link metal detoxification to immunity. Some studies suggest that there may be a connection between concentration of metal ions, such as iron, copper, cadmium and zinc, with plasmodesmatal permeability (O’Lexy *et al*., 2018), which might explain the function of a plasmodesmatal localized HIPP in regulating permeability, associated with their role in immunity. Both the HMA domain and isoprenylation regions of HIPPs have been shown to be important for plant immunity. For example, a mutation targeting the proline-rich region of *pi21* is sufficient to lead to gain of resistance against *M. oryzae* (Fukuoka *et al*., 2009), while interfering with the isoprenylation of HIPP1-V from wild wheat (*H. villosa*) leads to loss of resistance against *Blumeria graminis* f. sp. *tritici* accompanied by reduced HIPP1-V localization to plasma membrane. Interestingly, HIPP1-V interacts with the E3-ligase CMPG1-V at the plasma membrane leading to resistance to powdery mildew in an isoprenylation-dependent manner (Wang *et al*., 2023). This interaction has been reported to activate expression of genes involved in ROS generation and salicylic acid biosynthesis, suggesting that HIPP1-V is required for PTI regulation (Wang *et al*., 2023). Interestingly, we found that HIPP1-V is an ortholog of HvHIPP43 and can interact with Pwl2 in a Y2H assay. It is possible, therefore, that the Pwl2 interaction with HvHIPP43 attenuates PTI through a similar mechanism, which will require further investigation.

How the change in cellular localization of HIPP43 induced by the Pwl2 effector prevents its function in immunity or leads to a new role that enhances disease susceptibility remains unclear. A very recent study has provided evidence that *M. oryzae* AVR-Pik binding stabilizes the rice small HMA (sHMA) proteins OsHIPP19 and OsHIPP20 (Oikawa *et al*., 2024), suggesting that the function of Pwl2 may be mirrored by other blast effectors, targeting a wider pool of HIPPs. In this regard, future work will be necessary to determine whether HIPP43 directly regulates ROS generation, which might reduce permeability of plasmodesmata (Cui and Lee, 2016), or acts in a more indirect manner through interaction with other signaling components involved in PTI. Finally, Pwl2 is known to be a highly mobile effector and has been shown to move into neighbouring rice cells ahead of *M. oryzae* hyphal growth (Giraldo *et al*., 2013). This has been suggested to be a step to prepare un-invaded cells for fungal colonisation, consistent with its Pmk1-dependent regulation. Pwl2 may therefore alter the sub-cellular localisation and concentration of HIPP43, sequestering it away from PD that the fungus uses for effector movement and hyphal invasion, thereby enabling more rapid tissue colonisation by the blast fungus.

## Materials and methods

### Fungal strains, growth conditions and infection assays

Fungal isolates were routinely grown on complete medium (CM) at 24°C with a controlled 12 h light and dark cycle for up to 12 days (Talbot *et al*., 1993). *H. vulgare*, *E. curvula* and *O. sativa* plants were grown for 7, 14 and 21 days respectively in 9 cm diameter plastic plant pots or seed trays. Conidia were recovered from 10-day old cultures using a sterile disposable plastic spreader in 3 mL sterile distilled water. The conidial suspension was filtered through sterile Miracloth and centrifuged at 5000 x g for 5 min at room temperature before adjusting to a final of concentration of 1 x10^5^ conidia mL^-1^ in 0.2 % gelatin. The spore suspension was used for spray or leaf drop infections assays. After spray inoculation, plants were placed in polythene bags and incubated in a controlled plant growth chamber at 24°C for 48 h with a 12 h light-dark cycle and 85% relative humidity, before removing polythene bags. Inoculated plants were incubated for 3 days before scoring lesions. For each treatment 10 leaves were collected before counting typical ellipsoid necrotic disease lesions with a grey centre (Valent *et al*., 1991). Each experiment was repeated a minimum of three times, yielding consistent outcomes.

### Leaf infection assay and live-cell imaging

Rice leaf sheath from 4 week old susceptible cultivars Moukoto or CO39 were inoculated with 4 mL of a suspension at 5 x 10^4^ conidia mL^-1^ in ddH2O using a micropipette (Kankanala *et al*., 2007). Inoculated leaf sheaths were incubated at 24°C for 24 h before a thin layer of inner leaf sheath was dissected and mounted on a glass slide. Treatment with INA-PP1 was carried as described previously (Sakulkoo *et al*., 2018). Live cell imaging was carried out on an IX81 motorized inverted microscope (Olympus, Hamburg, Germany) for conventional and differential interference contrast (DIC) microscopy using Photometrics CoolSNAP HQ2 camera (Roper Scientific, Germany). Images were analyzed using ImageJ. For Leica SP8 laser confocal microscopy, settings were as follows; GFP, YFP and RFP/mCherry tagged proteins were excited using 488, 514 and 561 nm laser diodes and emitted fluorescence detected using 495-550, 525-565 and 570-620 nm respectively. Auto-fluorescence from chlorophyll was detected at 650-740 nm.

### Generation of fungal transformation plasmids

Single or multiple DNA fragments were cloned into fungal transformation vectors using In-Fusion HD Cloning (Clontech, USA). Briefly, fragments from cDNA, genomic DNA or synthesized DNA were amplified using primers to introduce a 15bp overhang complementary to sequences at restriction sites of a destination vector or adjacent insert fragments. This allows the ends to fuse by homologous recombination. Positive transformants were selected by colony PCR and constructs sequenced by GENEWIZ. A list of primers is provided in Table S3.

### RNA isolation, RNA sequencing and analysis

To study *in-planta* gene expression of *PWL2* and other effectors, leaf drop infection assays were carried out using susceptible rice cv. Moukoto and samples collected at 24 and 72 hpi. Infected plant material was ground to a fine powder using a sterile nuclease-free mortar & pestle containing liquid N_2_. RNA was isolated from *M. oryzae* mycelium, or inoculated rice leaves using QIAGEN RNeasy Plant Mini Kit. RNA quality was determined NanoDrop spectrophotometry (Thermo Scientific, UK) and Agilent 2100 Bioanalyser (Agilent Technologies, UK). Library preparation was carried out using Illumina® sequencing TruSeq Stranded Total RNA Library Prep Kit before sequencing 100 bp paired ends reads using Illumina Genome Analyser GXII platform by Exeter Sequencing Service (University of Exeter). To determine differential gene expression, raw reads were separated by mapping to both *M. oryzae* and *O. sativa* using kraken2. Reads specific to *M. oryzae* were used to quantify transcript abundance using Kallisto. To quantify genes missing in 70-15, separated reads were mapped to KE002. R package Sleuth was used to determine genes showing differential expression with log2fold > 1 and P-adjust value < 0.05 defined as up-regulated and a log2fold > 1 and P-adjust value < 0.05 as down-regulated. Southern blot analysis of *M. oryzae* genomic DNA was carried out as described previously (Talbot *et al*., 1993). Quantitative real-time PCR was carried out using CFX OPUS 96 and CT values normalised to a house keeping gene, β-Tubulin (MGG_00604). Fold change was determined using the formula 2-^ΔΔCT^, where *ΔΔ*Ct = ((CtGOI in infected sample - CtNC in infected sample) – (CtNC in mycelia - CtNC in mycelia)), and GOI is the gene of interest and NC (negative control) is β-Tubulin.

### Generation of Cas9-sgRNA targeted gene deletion

A sgRNA was designed for CRISPR-Cas9 genome editing using online tool E-CRISP http://www.e-crisp.org/E-CRISP/. A 20-nucleotide sequence was selected at the *PWL2* locus (not including the PAM NGG-sequence). The sgRNA was first synthesised using the EnGen sgRNA synthesis kit New England Biolabs (NEB #E3322) before mixing with Cas9-NLS to form an RNP complex (Foster *et al*., 2018). The mixture was incubated at room temperature for 10 min before fungal transformation. *M. oryzae* protoplasts were generated as described previously (Talbot *et al*., 1993). The RNP complex together with donor template was mixed with Guy11 protoplasts re-suspended in 150 µL STC to a concentration of 1×10^8^ mL^-1^ incubated at room temperature for 25 min before adding 60% PEG. Successful transformants were selected on complete medium (CM) agar containing 200 μg mL^-1^Hygromycin B.

### Whole genome sequencing

Purified RNA-free DNA was obtained using Hexadecyltrimethylammonium Bromide (CTAB). Template quality was assessed by NanoDrop and Qubit spectrophotometry. Sequencing was carried out at Exeter Sequencing services (University of Exeter, UK) and Novogene (Cambridge, UK). NEXTflex^TM^ Rapid DNA-seq Library Prep Kit was used to prepare and index libraries before sequencing on HiSeq 2500 (Illumina) with two lanes per sample. Quality of sequencing reads was checked using FastQC (http://www.bioinformatics.bbsrc.ac.uk/projects/fastqc/). From raw data (fastq files), adaptor sequences were trimmed from sequences containing adaptors and low-quality reads removed by fastq-mcf. Trimmed sequences were aligned to the reference genome (70-15) (Dean *et al*., 2005) using BWA (Burrow Wheeler Aligner) https://github.com/lh3/bwa <u>(Li and Durbin,</u> <u>2009)</u>. Bam files were visualized by IGV genome viewer to determine CRISPR gene deletion.

### Copy number variation of *PWL2*

A total of 286 *M. oryzae* isolates with raw Illumina-based sequencing information were downloaded from NCBI (performed on October 16^th^, 2019). Copy number variation was assessed using a k-mer analysis approach using the k-mer analysis toolkit (KAT; v2.4.1). Coding sequence information for *M. oryzae* isolate 70-15 was used as template for k-mer analysis and raw Illumina reads were input. Default parameters were used including k-mer length 27 nt. Copy number variation of individual effectors is based on average k-mer coverage compared with the median coverage for all genes.

### Phylogenetic analysis of *Magnaporthe* isolates

Genome sequences of diverse *M. oryzae* isolates were identified by literature review and searches on NCBI (performed August 9^th^, 2019). For isolates having only Illumina sequencing data, raw paired reads were downloaded from NCBI, trimmed using Trimmomatic (v0.36) using the parameters: removal of adapters with ILLUMINACLIP:TruSeq2-PE.fa:2:30:10, remove leading low quality or N based with quality below 5 for leading (LEADING:5) and trailing sequence (TRAILING:5), scan and cut reads with 4 bp sliding window below 10 (SLIDINGWINDOW:4:10), and minimum length 36 bp (MINLEN:36). KmerGenie (v1.7048) was used to identify an optimal k-mer for genome assembly using default parameters. Genome assembly was performed using minia (v0.0.102) with default parameters. kSNP3 (v.3.021) was used to develop a phylogenetic tree of *M. oryzae* using assembled genomes as input with parameters of k-mer of 29 bp and minimum fraction of 0.4. The phylogenetic tree was generated using RAxML (v8.2.12) using the General Time Reversible model of nucleotide substitution under the Gamma model of rate heterogeneity.

### Generation of *Agrobacterium* transformation plasmids

Single or multiple DNA fragments were cloned into binary vectors using In-Fusion HD Cloning (Clontech, USA). Briefly, fragments from cDNA were amplified to introduce a 15bp overhang complementary to sequences at restriction sites of a destination vector or adjacent insert fragments. Alternatively, synthesized DNA fragments were designed with a 15bp overhang complementary to sequences at restriction sites of binary vector pG514 customized for infusion cloning by digestion by either *Xba*I and *Pca*I or *Sac*I (New England Biolabs) to remove the *ccDB* toxin encoding gene. Positive transformants were accessed by colony PCR and constructs sequenced and analyzed on SNAP gene.

### Transient expression in *Nicotiana benthamiana*

*Agrobacterium tumefaciens* strain *GV3101* (Holsters *et al*., 1980) was used for transient expression. Three-week-old *N. benthamiana* leaves were infiltrated with transformed *Agrobacterium* carrying T-DNA constructs expressing the gene of interest. Bacterial cultures were diluted to obtain a final OD_600_ of 0.4 in agroinfilitration buffer (10mM MES, 10 mM MgCl_2_, 150 μM acetosyringone, pH 5.6). Leaf discs were cut from agroinfiltrated tissue 48 hpi and subjected for microscopy or used for co-IP.

### Plant transformation and ROS measurement

*Agrobacaterium tumefaciens* strain *AGL1* (Holsters *et al*., 1980) was used for plant transformation (Hensel *et al*., 2009). Positive transgenic plants were selected on HygromycinB followed by confirmation using PCR. Alternatively, leaf discs were collected from transgenic plants and analysed for copy number by iDNA Genetics Ltd (Norwich, UK). In addition, expression of the protein was evaluated using SDS-PAGE. To measure the response to PTI elicitors, a 4-mm diameter biopsy punch (Integra^TM^ Miltex ^TM^) was used to cut leaf discs from 5-week-old *H. vulgare* transgenic plants. Leaf discs were transferred to 96-well-plates (Greiner Bio-One) containing 100 µL ddH_2_O in each well and incubated overnight at room temperature. The assay was carried out by replacing ddH_2_O in wells by a 100 µL of 100 µM Luminol or L-012 (Merck), 20 µg mL^-1^ horseradish peroxidase (Merck) together with elicitors, flg22 (1 µM final concentration), chitin (1mg/mL final concentration) or without (mock). Photon count was carried out using a HRPCS218 (Photek) equipped with a 20 mm F1.8 EX DG ASPHERICAL RF WIDE LENS (Sigma Corp). Each experiment was repeated three times, yielding consistent outcomes.

### Co-immunoprecipitation and sample preparation for mass spectrometry

Leaves were harvested from 14-21 day old barley transgenic plants and rapidly frozen in liquid N_2_ before storage or immediately ground to fine powder using a GenoGrinder® tissue homogenizer. Ground powder was quickly transferred to ice-cold 1.5 mL Eppendorf tubes and 2 mL of ice-cold extraction buffer (GTEN [10% (v/v) glycerol, 25 Mm Tris pH 7.5, 1 mM EDTA 150 mM NaCl, 2% (w/v) PVPP, 10 Mm DTT, 1 X protease inhibitor cocktail (Sigma), 0.5% (v/v) IGEPAL added and mixed thoroughly. This was centrifuged at 3000 x g for 10 min at 4°C to recover total protein in the supernatant and repeated twice. A volume of 2 mL of total protein was mixed with 25 μL of GFP-TRAP agarose beads (50% slurry, ChromoTek) and incubated shaking overnight at 4°C. The GFP-TRAP agarose beads were centrifuged for 2 min at 3000 x g before washing three times with wash buffer (ChromoTek). Bound proteins were recovered by resuspending beads in loading buffer (loading dye, 10 mM DTT, H2O) before incubating at 70°C for 10 min. An aliquot was run by SDS-PAGE and used for immuno-blot analysis (Burnette, 1981). Recovered proteins were fractionated by SDS-PAGE for approximately 1 cm. The gel was washed for 1 h in ddH_2_O followed by 1 incubation SimplyBlue^TM^ Safe stain. The gel was then washed three times in ddH_2_O and the region containing proteins excised using a scalpel.

### Mass spectrometry and data processing

Protein purification, immunoprecipitation, sample preparation, liquid chromatography followed by tandem mass spectrometry (LC-MS/MS), and data analysis were carried out as previously described (Li *et al*., 2023). Proteins identified in immunoaffinity-enriched samples were measured by high resolution LC-MS systems, Orbitrap Fusion (Thermo Ltd). Acquired spectra were peak-picked and searched by Mascot (Matrix Science Ltd) to identify peptide sequences from the search space defined by background proteome. Peptides were combined into proteins based on the principle of parsimony by the search engine. Resulting proteins were further described by quantitative values based on the number of spectra that identify them. Individual runs were combined in Scaffold (Proteome Software Inc.), where the data were evaluated and filtered to contain less than 1% false positives (FDR) and the resulting matrix exported. The matrix of proteins detected in different samples serves as the input for an R script for further processing and visualization.

### Yeast two-hybrid analysis

To clone genes of interest into bait (pGBKT7 DNA-BD cloning) vector or prey (pGADT7 activating domain) vector, by In-Fusion HD Cloning (Clontech), bait and prey vectors were digested by *Bam*H1 and *Eco*R1 while gene to be inserted was PCR-amplified using primers containing 15 bp overhangs with homology to two ends of the digested bait vector. For transformation, a single colony Y2H gold yeast strain was mixed in 1 mL of liquid yeast extract peptone dextrose (YPD). Competent cells were prepared according to manufacturer instructions (Zymo Research). Briefly, for transformation 700 ng – 1 µg of plasmids expressing Pwl2 or other effectors in pGADT7 and *Hv*HIPP43 or plant proteins in pGBKT7 were co-transformed in competent cells and Frozen-EZ Yeast Solution 3 added before incubation at 28°C for 1 h and transformed cells plated on selection media lacking Leucine (L) and Tryptophan (W) and incubated at 28° C for 3-5 days. To detect interactions colonies were transferred to media lacking Leucine (L), Tryptophan (W), Adenine (A) Histidine (H), Bromo-4-Chloro-3-Indolyl a-D-galactopyranoside (X-a-gal) and 10 mM 3-amino-1,2,4-triazole (3AT) (Sigma). The plates were imaged after 60-72 incubation at 28°C. Each experiment was conducted a minimum of three times.

### *In-planta* co-immunoprecipitation (co-IP)

To test interactions of proteins *in planta*, genes of interest tagged with either GFP or Myc were cloned into vector pGW514, transformed into *Agrobacterium* strain *GV3101* and co-infiltrated (OD_600_ = 0.4 for effectors and OD_600_ = 0.6 for HIPP43) into 3-4 week-old *N. benthamiana* leaves and incubated for 48 h to allow protein expression. Total protein was isolated in ice-cold extraction buffer (GTEN [10% (v/v) glycerol, 25 Mm Tris pH 7.5, 1 mM EDTA 150 mM NaCl], 2% (w/v) PVPP, 10 Mm DTT, 1 X protease inhibitor cocktail (Sigma), 0.5% (v/v) IGEPAL). Total protein was co-immunoprecipitated using anti-GFP M2 resin (Sigma-Aldrich, St. Louis, MO) and washed 3 times using immunoprecipitation buffer before analyzing by SDS-PAGE. Recovered proteins from co-immunoprecipitation were separated by SDS-PAGE and transferred to a polyvinylidene diflouride (PVDF) membrane using a Trans-Blot turbo transfer system (Bio841 Rad, Germany). Detection was performed using the appropriate antibody either anti-GFP-HRP or anti-Myc-HRP. Imaging was carried out using an ImageQuant LAS 4000 luminescent imager (GE Healthcare 844 Life Sciences, Piscataway, NJ, U.S.A.) according to manufacturer’s instructions.

### Sub-cellular localization and plasmodesmata quantification

Leaf discs were obtained from *N. benthamiana* plants agroinfiltrated with different construct combinations at 24 and 48 hpi and mounted on a slide immersed in perfluorodecalin and observed using X63 oil immersion lens. For Leica SP8 laser confocal microscopy, settings were as follows; GFP, YFP and RFP/mCherry tagged proteins were excited using 488, 514, and 561 nm laser diodes and emitted fluorescence detected using 495-550, 525-565, and 570-620 nm respectively. Auto-fluorescence from chlorophyll was detected at 650-740 nm. To stain and quantify plasmodesmata (PD), agroinfiltrated leaves were further infiltrated with 0.1% aniline (Sigma-Aldrich #415049, in PBS buffer, pH 7). Images were analyzed using ImageJ to determine the number of PD. First, cell peripheries were divided into 20 µm sections and the number of PD determined per 20 µm.

### Phylogenetic analysis of grass HMA proteins

Proteins from *B. distachyon* (314; v3.1), *H. vulgare* (Barley cv. Morex V3, Jul 2020), *T. aestivum* (RefSeqv2.1; 09-16-2020), *O. sativa* (323; v7.0), *Oropetium thomaeum* (386; v1.0), *Sorghum bicolor* (454; v3.1.1), *Setaria italica* (312; v2.2), and *Zea mays* (RefGen_V4) containing an HMA domain were identified using InterProScan (v5.59-91.0; Pfam PF00403). The HMA domain was extracted using the script QKdomain_process.py (https://github.com/matthewmoscou/QKdomain) including an additional 10 amino acid sequences N and C-terminal of the Pfam boundaries (-n 10-c 10). The non-redundant set of HMA domains were identified using CD-HIT (v.4.8.1) with parameter-c 1.0. Structure-guided multiple sequence alignment was performed using MAFFT with parameters dash, max iteration of 1000, and globalpair. Maximum likelihood phylogenetic analysis was performed using RAxML (v8.2.12) using the Gamma model of rate heterogeneity, JTT amino acid substitution model,-and 1,000 bootstraps. Coding sequence was identified for the HIPP43 gene family and aligned using MUSCLE (v5) translation alignment using default parameters. Maximum likelihood phylogenetic analysis was performed using RAxML (v8.2.12) using General Time Reversible model of nucleotide substitution under the Gamma model of rate heterogeneity and 1,000 bootstraps.

### Statistical analysis and Protein Structure Prediction

Significance difference between groups of samples was performed using GraphPad Prism 10. *P* values < 0.05 were considered significant, \*\*\*\**P*<0.0001, \*\*\**P*<0.001, \*\**P*<0.01 while values *P* values > 0.05 were considered as non-significant in an unpaired Student’s t-test. Structure prediction was carried out using AlphaFold3 (Abramson *et al*., 2024), the structure analyzed and figures generated using ChimeraX (Meng *et al*., 2023).

## Supporting information

Supplemental Data 1. Blast genomes used for phylogenetic analysis

Supplemental Data 4. IP-MS samples report with clusters

Supplemental Data 3. IP-MS spectrum report

Supplemental Data 2. Blast genomes used for copy number variation analysis.

## Acknowledgements

We thank current and past members of the “BLASTOFF” team from the Banfield, Kamoun, Terauchi, Talbot, and Moscou Laboratories. For funding, we thank Gatsby Charitable Foundation and Biotechnology and Biological Sciences Research Council (BBSRC) BBS/E/J/000PR9797 and a BBSRC grant to NJT and FLHM (BB/V016342/1), United States Department of Agriculture-Agricultural Research Service CRIS #5062-21220-025-000D (MJM) and BB/X010996/1, funding for APH ISP. We acknowledge Y.K Gupta, J.C De la Concepcion, J. Rhodes and J. Win for constructive discussions.

## Author Contributions

**Conceptualization:** NJT and VW

**Formal analysis:** VW, XY, AF, JK, JS, DG, TL, YPH, ABE, MS, MJB, MM, WM and RZ

**Funding acquisition:** NJT

**Investigation:** VW, XY, AF, JS, DG, AG, NS, DK, YPH, ABE, MJB, WM and RZ

**Writing – original draft:** VW and NJT

**Writing – review & editing:** VW, NJT, LR, AB, TL, MB, SK, MM, FM

---

**Fig. S1.**
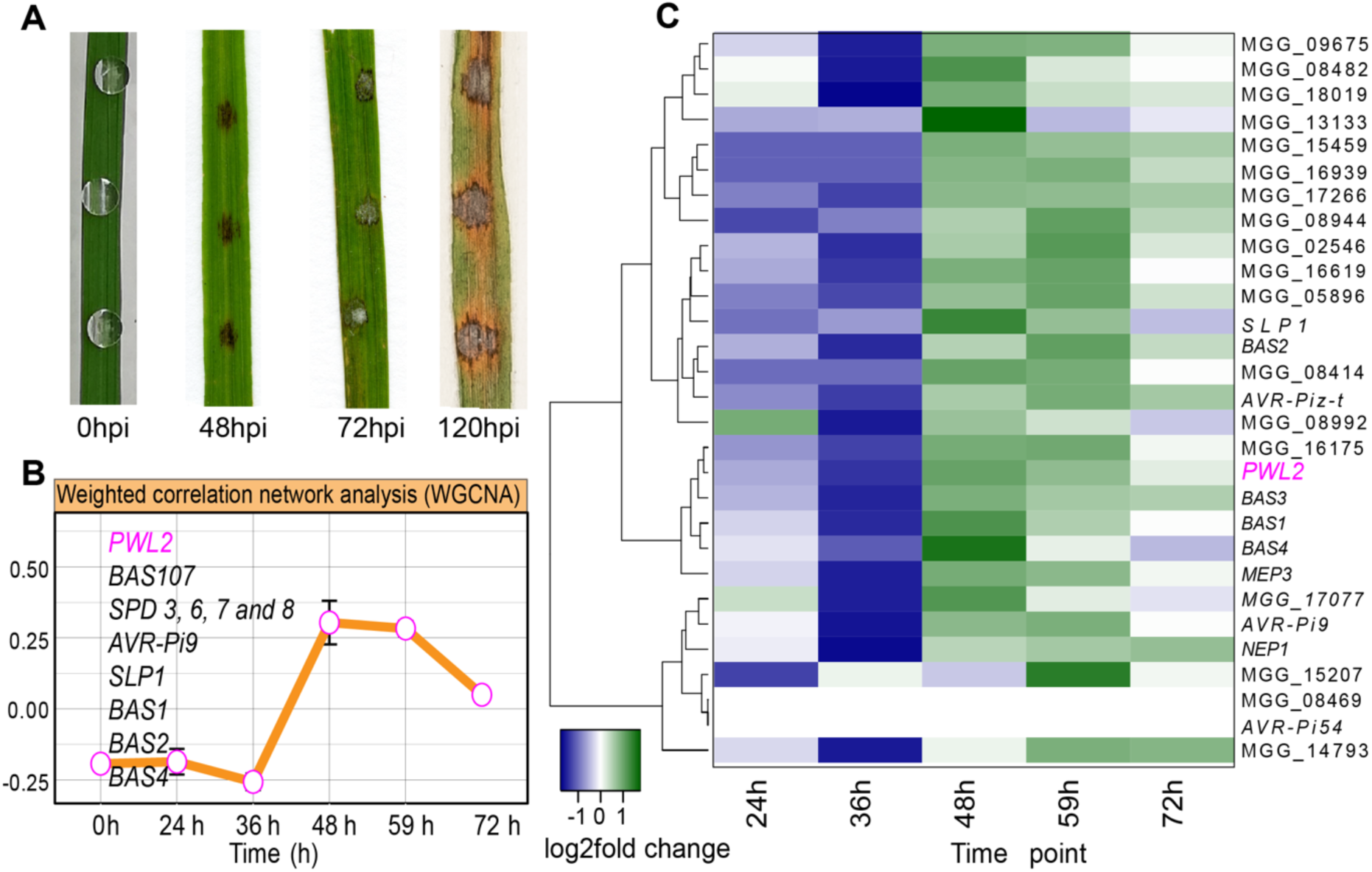
*M. oryzae* Pwl2 is co-expressed during plant infection with other known effector genes. (**A**) Attached leaves of three-weeks old rice seedlings inoculated with KE002 conidia suspension at 0, 48, 72 and 120 hpi. Samples were collected at 24, 36, 48, 59 and 72 hpi and used for RNA isolation (3 biological replicates). (**B**) RNA isolated from infected rice leaves were sequenced and the reads normalized against *M. oryzae* mycelia KE002 RNA to determine differentially expressed genes. A weighted gene co-expression network analysis grouped gene expression profile into 10 co-expression modules. The line graph (WGCNA) shows the representative eigengene for module 4 in which *PWL2* is co-expressed with other known effector protein encoding genes, *AVR-Pi9*, *SLP1*, *BAS1-4* and suppressors of cell death effectors 3, 6, 7 and 8 that possibly serve related function during biotrophic phase of invasion. (**C**) Heatmap showing relative expression levels (log2fold change) of *PWL2* and other known effectors during infection.

**Fig. S2.**
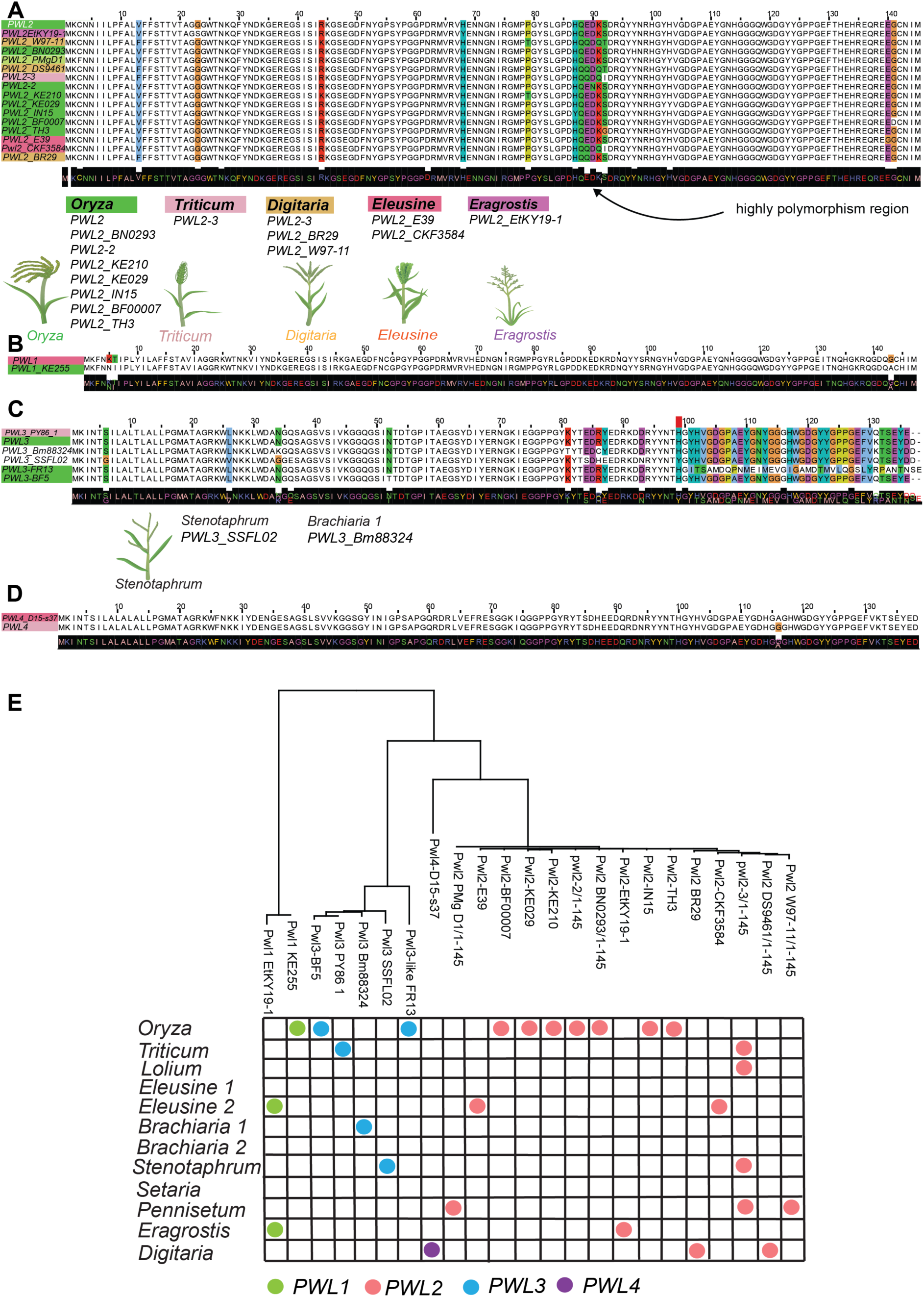
***PWL2* is highly polymorphic compared to other members in the *PWL* family.** (**A**) Multiple amino acid sequence alignment of Pwl2 coding sequence and naturally occurring alleles. The region between His-87 and Ser-92 shows highest number of polymorphisms and is the region important for cognate resistance gene recognition. Multiple amino acid sequence alignment of *PWL1* (**B**), *PWL3* (**C**) and *PWL4* (**D)** Alleles identified from genomes of field isolates. Alignments were carried using ClustalW (Jalview). (**E**) Neighbor-joining tree based on amino acid sequence of alleles of the *PWL* gene family. Tree IDs contains isolate name from which allele was identified, and each dot represents presence in each host-limited form. Unlike other family members, *PWL2* alleles are present in host-limited forms apart from lineage 2 of *Brachiaria* and *Setaria. Panicum, Cynodon* and *Urochloa*-infecting isolates.

**Fig. S3.**
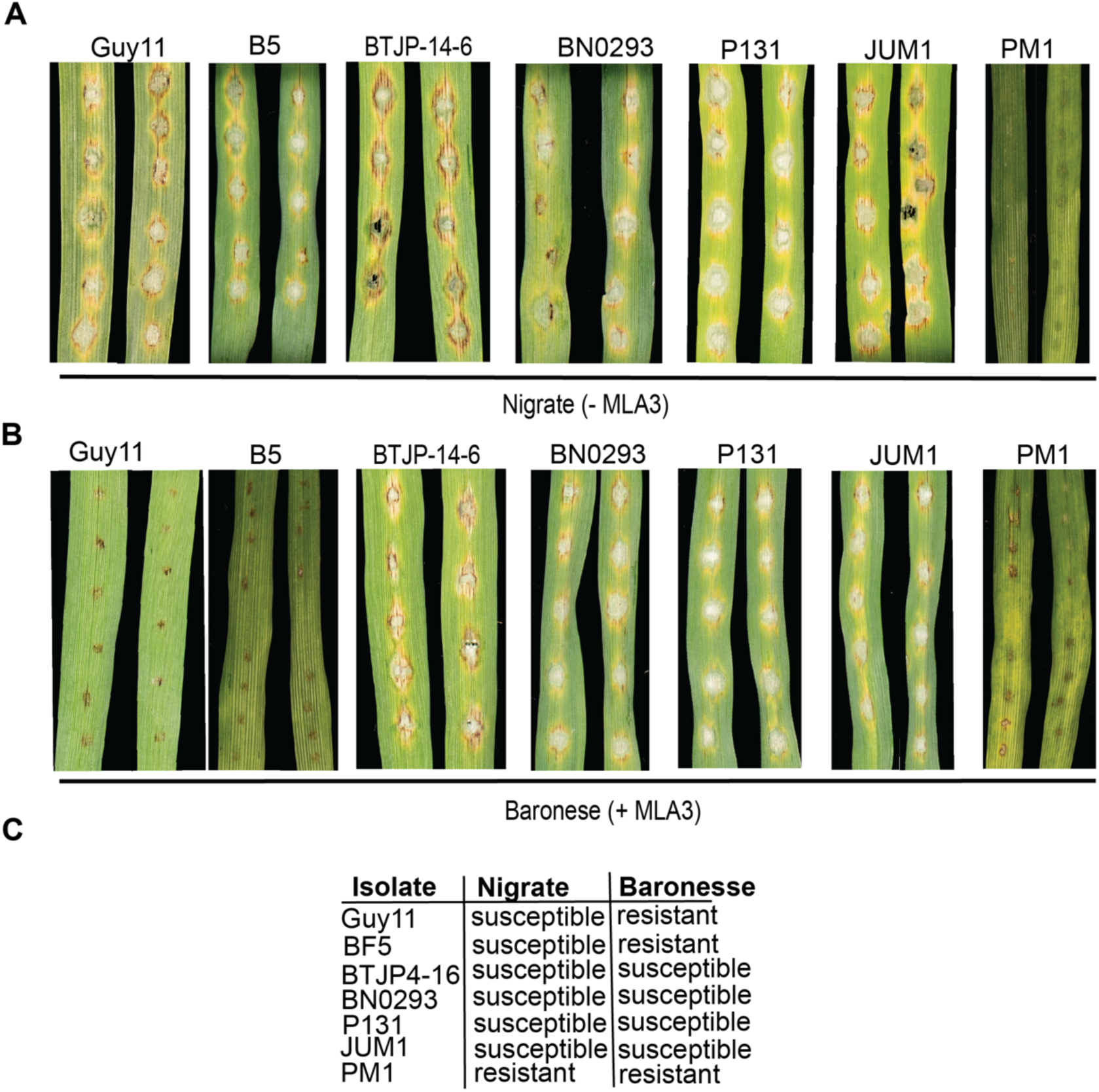
Two *PWL2* alleles are not recognized by the barley resistance gene Mla3. A subset of isolates carrying alleles of *PWL2* were used in leaf drop inoculation on (**A**) barley accessions cv. Nigrate (-*Mla3*) and (**B**) cv. Baronesse (+*Mla3*). Conidial suspensions at 1 ×10^5^ mL^−1^ spores/mL were harvested from Guy11 and selected isolates with *PWL2* alleles and used to inoculate 10-day-old seedlings of cv. Nigrate (-*Mla3*) and cv. Baronesse (+*Mla3*). Disease lesions were scored 5 days post-infection. Isolates BF5 (variant *Pwl2*), BTJP4-16 (*pwl2-3)*, BN0293 (variant *Pwl2*), PM1 (variant *Pwl2*), Guy11 (+*PWL2*), JUM1 (*pwl2-2*), and P131 (*-PWL2*) were used for leaf drop infection assay. As expected Guy11 could infect cv. Nigrate but not cv. Baronesse. BF5 and PM1 did not produce blast disease on barley cv. Baronesse, suggesting they express *PWL2* alleles recognised by Mla3, (or express effectors recognised by additional R-genes contained (in barley cv. Baronesse). Notably, PM1 could also not infect cv. Nigrate (-*Mla3*). P131 does not carry *PWL2* and caused blast disease on cv. Baronesse as expected. JUM1 (*pwl2-2*), BTJP4-16 (*pwl2-3)* and BN0293 (*pwl2-BN0293*) all caused blast disease on cv. Baronesse (**C**) Summary of disease reaction of isolates with distinct *PWL2* alleles on Nigrate (-*Mla3*) and Baronesse (+*Mla3*). Observations were consistent in three independent infection replicates.

**Fig. S4.**
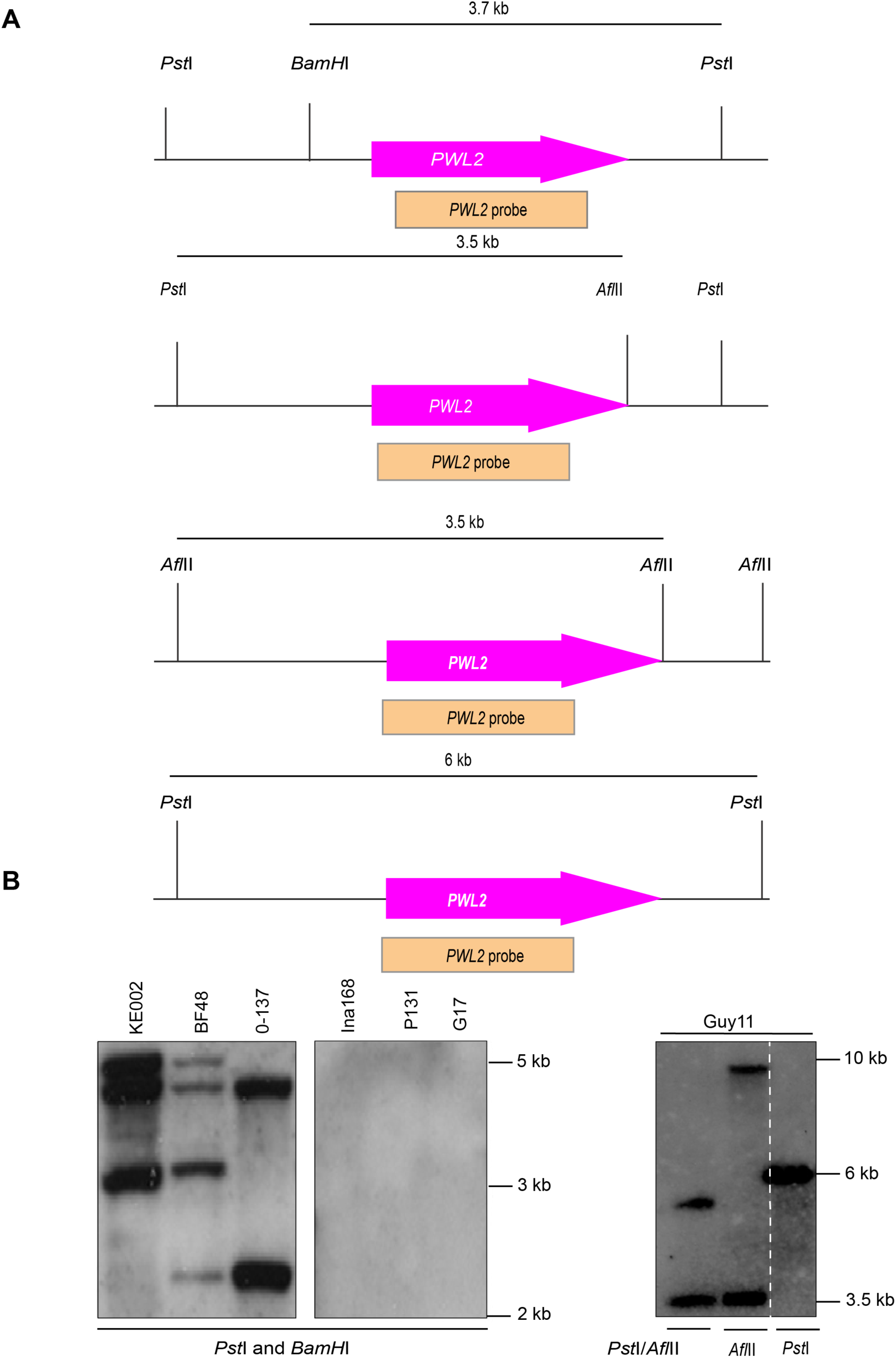
Southern blot analysis to determine copy number variation of *PWL2* in selected ***M. oryzae* isolates.** (**A**) Genomic DNA was digested using different enzyme combination *Bam*HI/*Pst*I (upper panel), *Pst*I/*Afl*II (upper middle panel), *Afl*II single digest (lower middle panel) or *Pst*I single digest (lower panel) and probed using the *PWL2* 438 bp probe. Size estimates are from the *PWL2* loci annotated *M. oryzae* 70-15 genome. (**B**) Left panel, the order of digested genomic DNA from different isolates is indicated on the lanes. Right panel, when Guy11 genomic DNA was digested using *Pst*1 and *Afl*II, we detected two bands of 3.5kb and 6kb. When digested with *AflII in a* single digest were detected two bands of 3.5 kb and 10 kb. While in a single digest using *Pst*I only produce a single band of 6.5 kb.

**Fig. S5.**
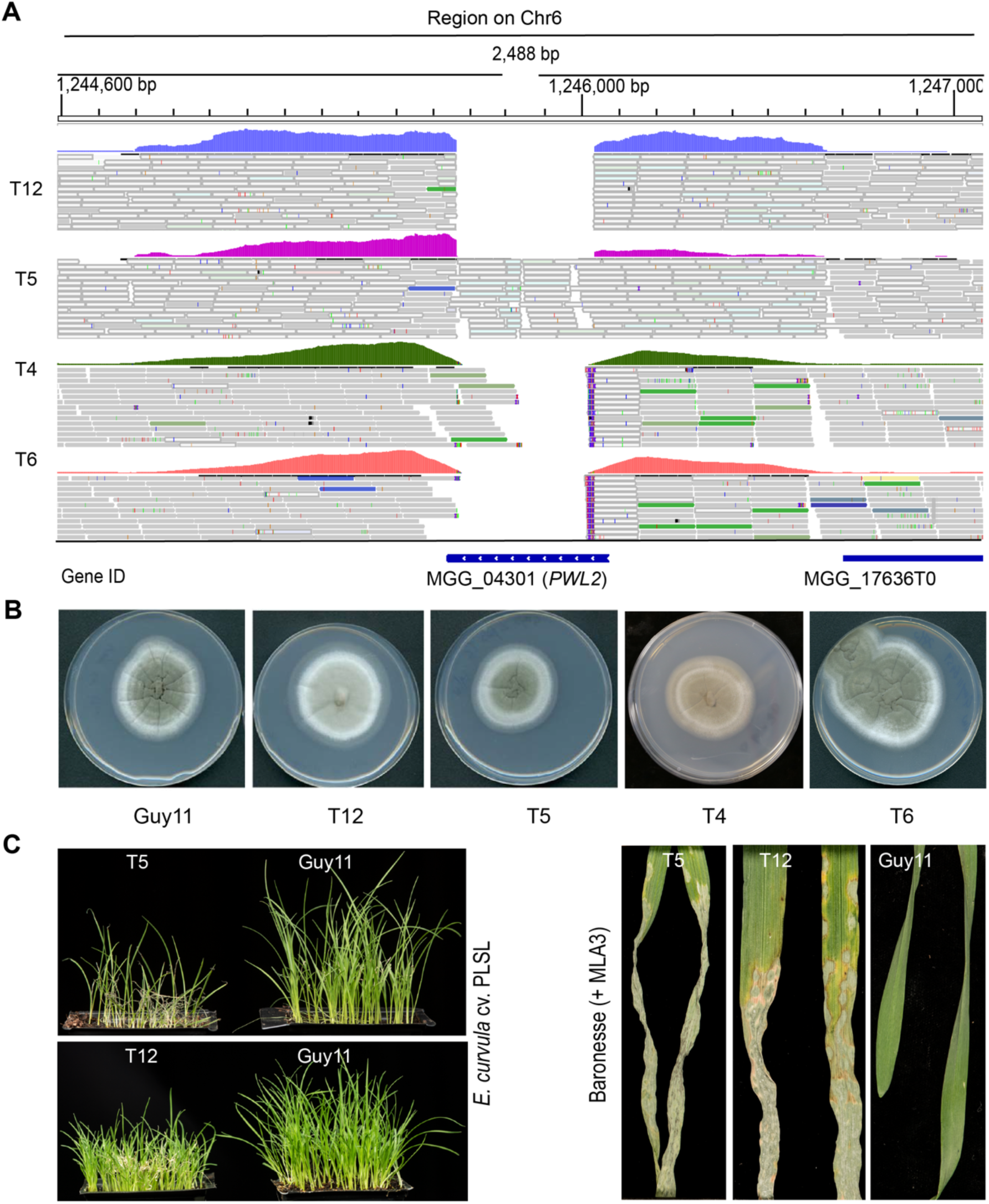
*M. oryzae Δpwl2* deletion mutants show gain of virulence on weeping lovegrass and cultivar Baronesse (*Mla3*). (**A**) Reads alignment display of *PWL2* locus on Chromosome 6 from four sequenced *pwl2* mutant strains, T12, T5, T4 and T6 (lower panel). No reads aligned to the MGG_04301T0 (*PWL2*) locus for T6 and T12 and partial alignment for T4 and T5. Read depth around *PWL2* locus is higher compared to the neighbouring gene MGG_17636T0 suggesting the *PWL2* locus occurs multiple times. Read coverage and locus length are indicated. (**B)** Guy11, T12, T5, T4, T6 were grown at 25^0^ C, on CM media for 10 days before imaging. (**C)** Disease symptoms on weeping lovegrass (*E. curvula*) and barley cultivar Baronesse (*Mla3*) to show gain of virulence for CRISPR-Cas9 deletion mutants T5 and T12. Conidial suspensions at 1 ×10^5^ mL^−1^ spores/mL from Guy11, *Δpwl2* (T5) and *Δpwl2* (T12) were used to inoculate 10-day-old seedlings of barley cultivar Baronesse (*Mla3*) or weeping lovegrass, and disease symptoms recorded after 5 dpi. Observations were consistent in three independent infection replicates.

**Fig. S6.**
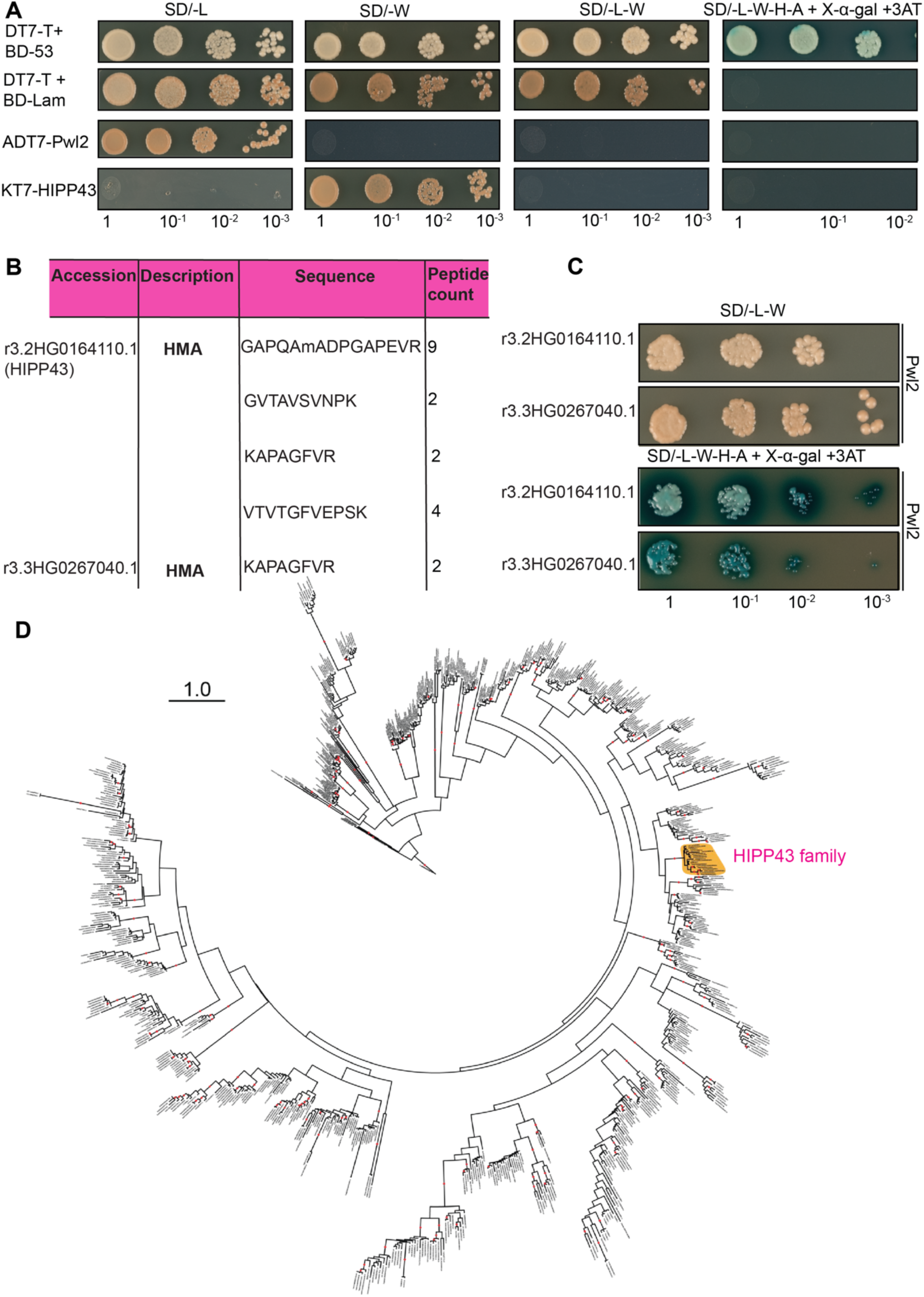
Pwl2 interacts with two copies of *Hv*HIPP43 identified by IP-MS. **(A**) Y2H assay to show that Pwl2 (in pGADT7 vector) and HIPP43 (in pGBKT7 vector) do not show autoactivity (**B**) Table showing peptides identified by LC-MS that mapped to two copies of HIPP43. (**C**) Pwl2 interacts with the two copies of HIPP43 in a yeast-two-hybrid assay. (**D**) Maximum likelihood phylogenetic tree of HMA domains from diverse grass species. HMA domains were identified from proteins from the grass species *B. distachyon*, *H. vulgare*, *T. aestivum*, *O. sativa*, *Oropetium thomaeum*, *Sorghum bicolor*, *Setaria italica*, and *Zea mays*. Structure guided multiple sequence alignment was performed using MAFFT-DASH and phylogenetic analysis with RAxML (v8.2.12). Scale bar indicates substitutions per site. Dots on branches indicate support of 80% or more based on 1,000 bootstraps. The HIPP43 family formed a distinct clade with bootstrap support of 98%.

**Fig. S7.**
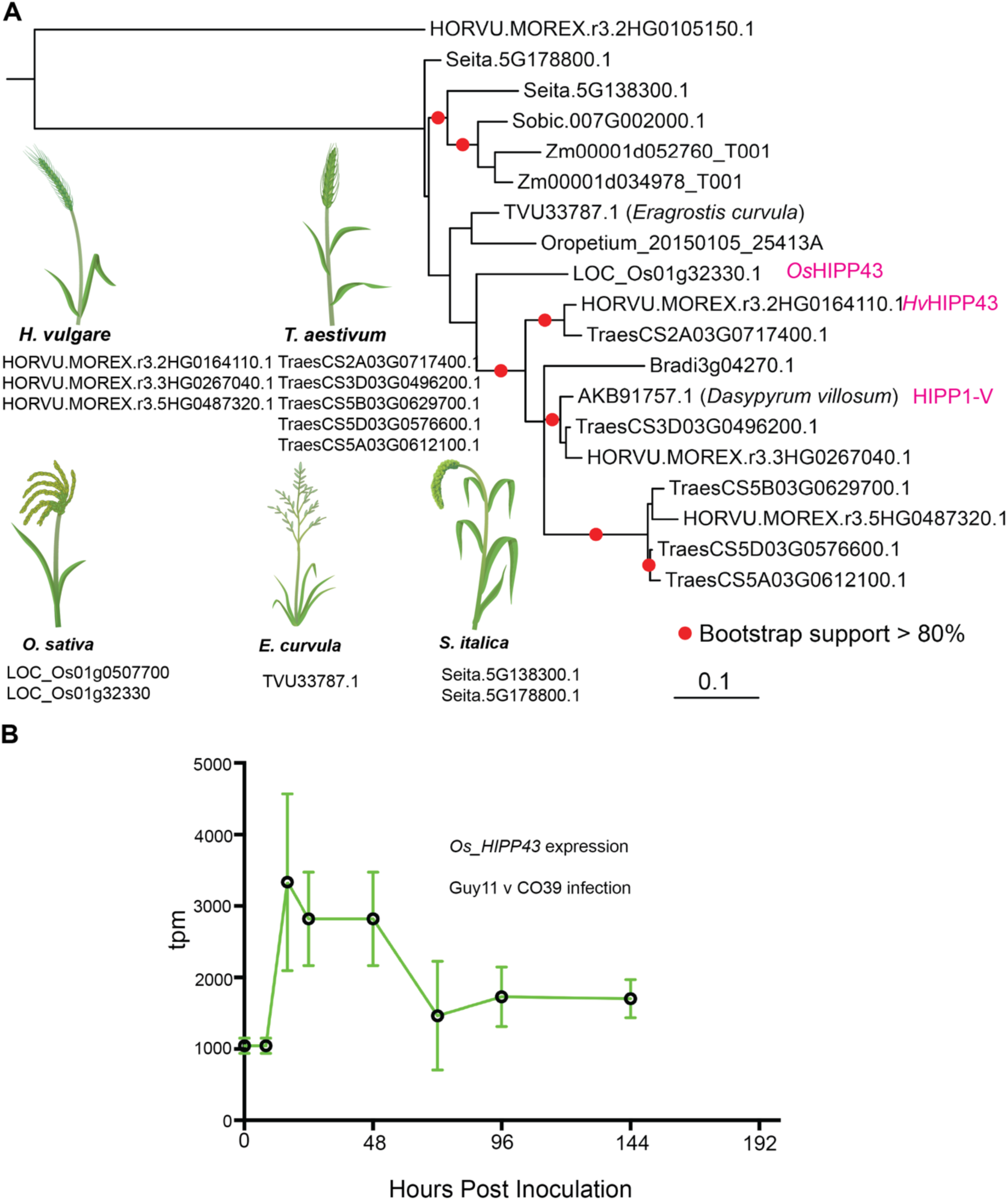
Phylogenetic analysis of HIPP43 and expression profile during the rice blast infection on a susceptible rice cv. CO39. (**A**) Full length coding sequence was identified for HIPP43 gene family members, aligned using MUSCLE translation alignment, and phylogenetic analysis with RAxML (v8.2.12). Barley and wheat (i.e. *Triticeae* lineage) have three copies of HIPP43 per haploid genome. Dots on branches indicate support of 80% or more based on 1,000 bootstraps. (**B**) RNA-seq analysis of *OsHIPP43* expression during infection of a susceptible rice cv. CO39 by Guy11. Hour post inoculation is shown on the X –axis. HIPP43 expression is shown as relative mean expression TPM (transcripts per million) at different stages of infection.

**Fig. S8.**
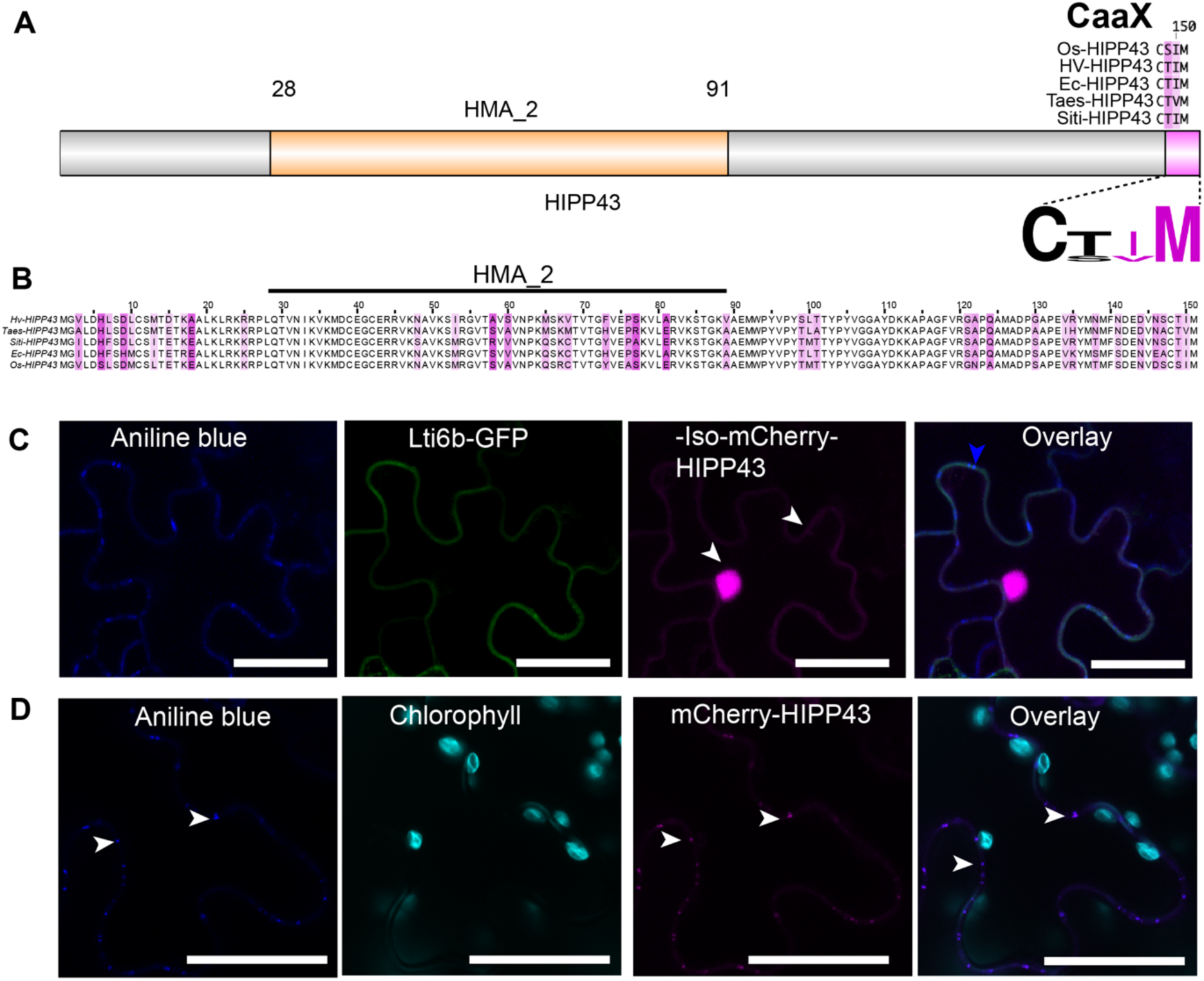
HIPP43 isoprenylation motif is required for PD localisation. (**A +B**) A schematic representation of domain structure of HIPP43 showing the HMA domain region and the isoprenylation motif at the C-terminus. Residues occurring at the C-terminus of HIPP43 orthologs in rice (Os-HIPP43), wheat (Taes-HIPP43), foxtail millet (Siti-HIPP43) and weeping lovegrass (Ec-HIPP43) are indicated. A multiple alignment was generated using Clustal X in Jalview and the conserved motif visualized on Weblogo. (**C-D**) Micrographs showing localization of HIPP43 without the isoprenylation motif (-Iso-mCherry-HIPP43) and of WT mCherry-HIPP43. (**C**) Deletion of the isoprenylation abolishes PD localisation but HIPP43 localises in the nucleus, the plasma membrane after 48h. Co-expression with a plasma membrane marker LTi6b-GFP and aniline blue staining confirmed localisation at the plasma membrane and lack of PD localisation. White arrows indicate regions of HIPP43 localisation away from aniline blue stained PD **(D)** Micrographs showing typical mCherry-HIPP43 localizing as small puncta on the plasma membrane when expressed in *N. benthamiana.* Staining of callose using aniline blue overlaps with mCherry-HIPP43, confirming mCherry-HIPP43 localises exclusively at PD. White arrows indicate regions of co-localisation between HIPP43, and aniline blue stained PD. Scale bars represent 20 µm.

**Fig. S9.**
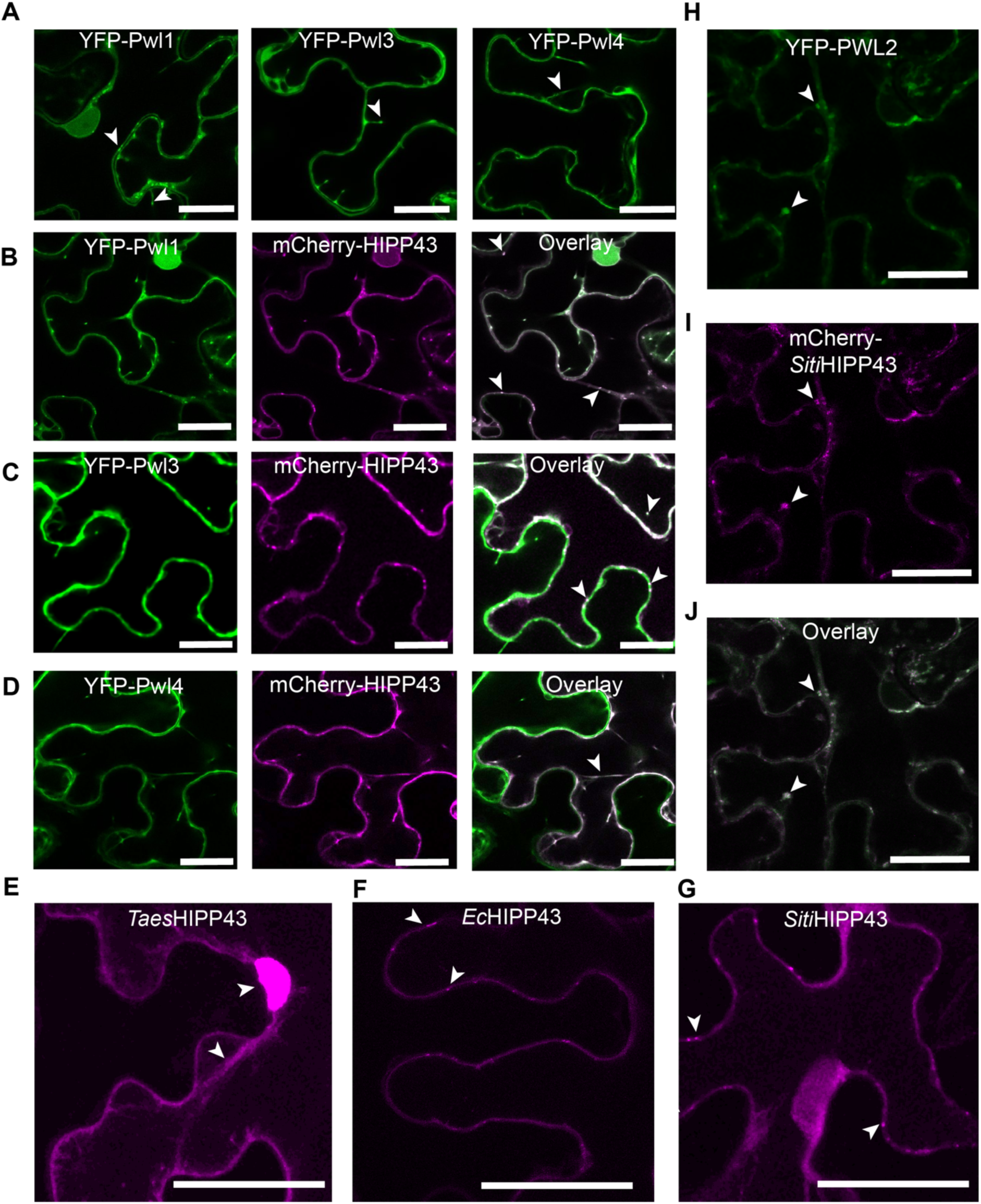
Pwl1, Pwl3, Pwl4 co-localise with HIPP43 and Pwl2 co-localises with HIPP43 orthologs from other grass species. (**A**) Micrographs showing localization of YFP-Pwl1, YFP-Pwl3 and YFP-Pwl4. Like YFP-Pwl2, Pwl1, Pwl3 and Pwl4 showed localisation in cytoplasm and cytoplasmic mobile structures. **(B-D)** Micrographs showing co-localisation of Pwl1, Pwl3 and Pwl4 with m-Cherry-HIPP43. **(B)** In presence of Pwl1 mCherry-HIPP43 translocates to the cytoplasm and partially in cytoplasmic mobile structures. **(C)** Pwl3 did not dislodge mCherry-HIPP43 from the PD while Pwl4 **(D)** only showed cytoplasmic co-localisation. (**E-G**) Micrographs showing HIPP43 orthologs fluorescently tagged to mCherry. (**E**) *Taes*HIPP43 (wheat) localizes in the nucleus and cytoplasm while (**F**) *Ec*HIPP43 (weeping lovegrass) (**G**) *Siti*HIPP43 (foxtail millet) localize as small puncta equivalent to PD. Images shown as maximum projections **(H-J)** Micrographs showing co-localisation of YFP-Pwl2 with mCherry-*Siti*HIPP43. mCherry-*Siti*HIPP43 localisation, was altered to the cytoplasm, cytoplasmic mobile structures. mCherry-*Siti*HIPP43 fluorescence signal was higher in presence of YFP-Pwl2. White arrows indicate regions of co-localisation and blue arrows show aniline blue stained PD. Scale bars represent 20 µm.

**Fig. S10.**
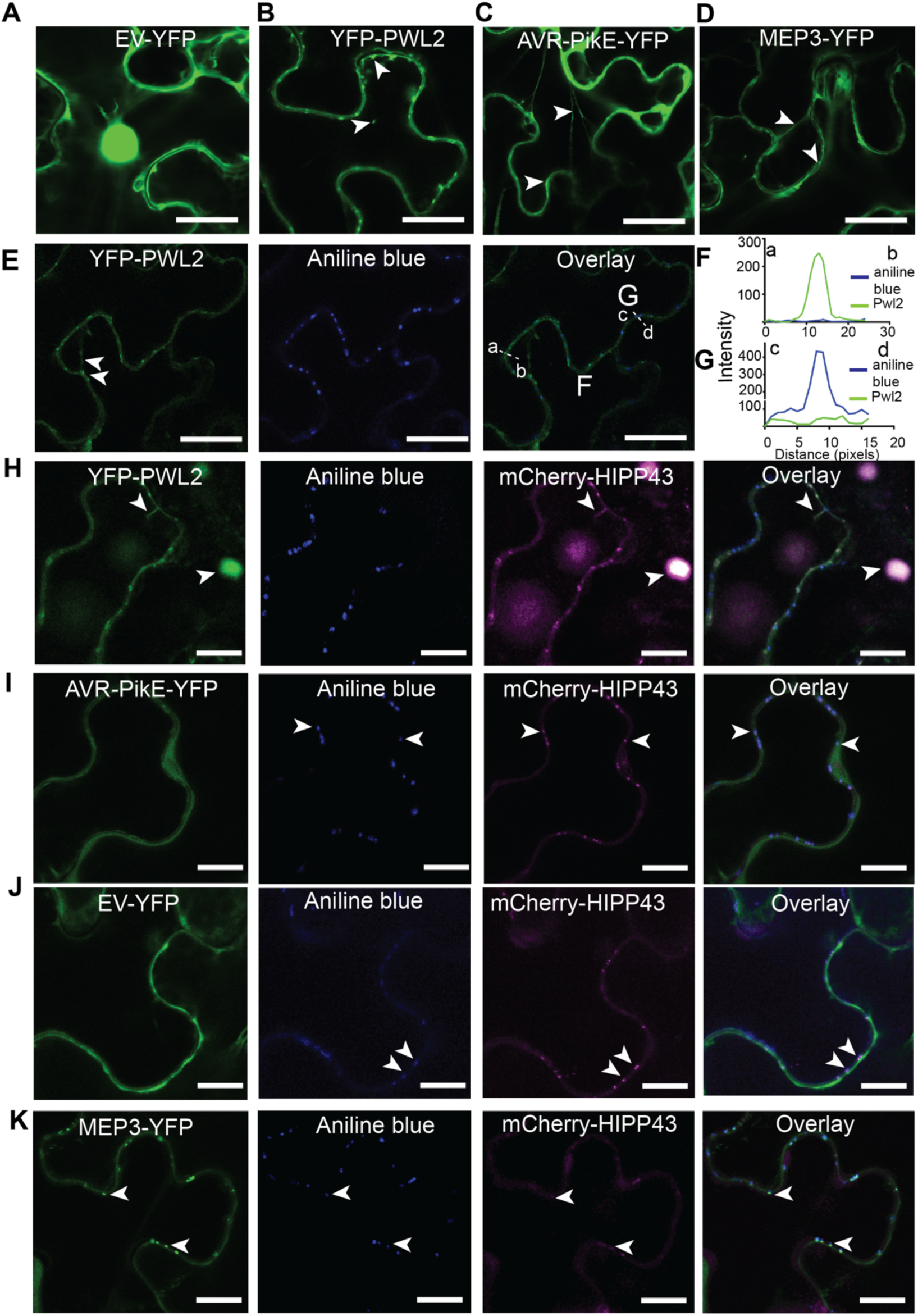
Pwl2 stabilises and promotes HIPP43 cytoplasmic localisation. (**A-D)** Micrographs showing localization of free-YFP (**A**), AVR-PikE-YFP (**B**), YFP-Pwl2 (**C**) and MEP3-YFP (**D**). Both free-YFP and AVR-PikE-YFP show localisation in the cytoplasm while YFP-Pwl2 showed localisation in cytoplasm and cytoplasmic mobile structures. MEP3 localises in the cytosol and PD **(E-I**) Micrographs and line scan graphs showing the presence of YFP-Pwl2 in the cytosol and puncta. Lack of co-localization with aniline blue stain confirms that, without HIPP43, Pwl2 does not accumulate at PD but rather mobile puncta. (**H-K**) Micrographs showing changes of mCherry-HIPP43 localisation when co-expressed with (**H**) YFP-Pwl2, (**I**) AVR-PikE-YFP (**J**) EV-YFP and (**K**) MEP3-YFP. Staining of callose using aniline blue is used to differentiate between immobile PD and mobile cytoplasmic mCherry-HIPP43 co-localisation with YFP-Pwl2. In presence of AVR-PikE, EV and MEP3 aniline blue staining co-localises with immobile mCherry-HIPP43. Scale bars represent 20 µm.

**Fig. S11.**
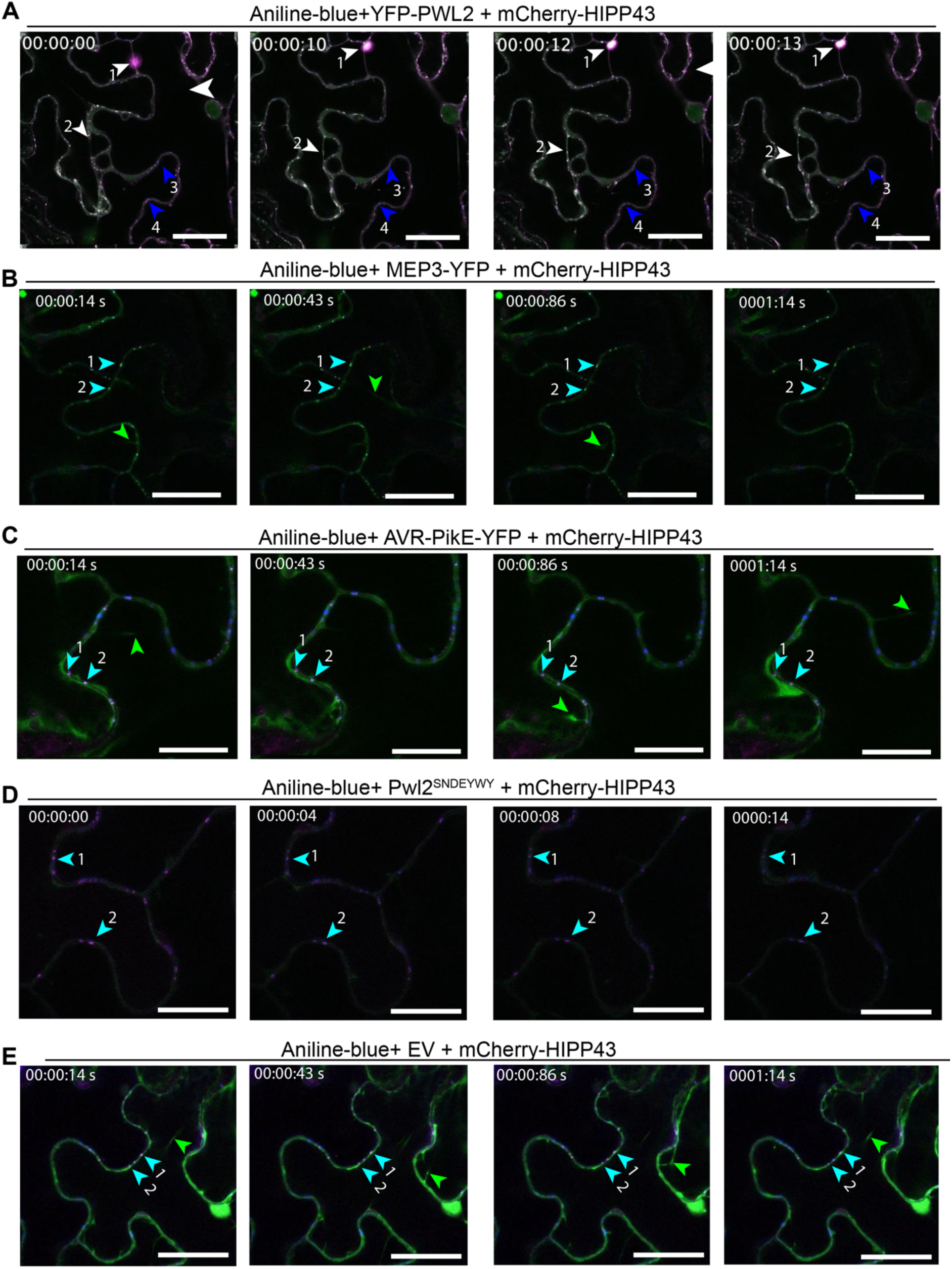
Pwl2 alters HIPP43 cellular localisation from the plasmodesmata. (**A**) A time-lapse confocal microscopy micrograph to shows changes of mCherry-HIPP43 localisation upon interaction with Pwl2 after co-expression in *N. benthamiana.* Pwl2 and HIPP43 co-localise as small or large mobile puncta. White arrow heads indicate mobile Pwl2/HIPP43 co-localisation puncta and blue arrow heads show immobile aniline blue stained PD. Arrow 1 indicate large cytoplasmic mobile body, arrow 2 show small mobile puncta. (**B-E**) Shows lack of co-localisation between MEP3 (**B**), AVR-PikE (**C**), Pwl2^SNDEYWY^ (**D**) and (**E**) free-YFP with HIPP43. HIPP43 co-localise with PD aniline blue stain as shown with cyan arrow heads and remains immobile. Note, MEP3 localises at PD in presence or absence of HIPP43. Green arrows show expression of AVR-PikE, MEP3 and free-YFP in the cytosol without HIPP43 co-localisation. Scale bars represent 20 µm. Aniline blue is shown in blue, free-YFP, Pwl2, MEP3, AVR-PikE and Pwl2^SNDEYWY^ are shown as green while mCherry-HIPP43 is shown as magenta.

## List of Movies

**Movie S1**

Mobility of Pwl2/HIPP43 sub-cellular co-localisation cytosolic puncta, 48 h after co-expression in *N. benthamiana* cells and visualized using laser confocal microscopy. PD are visualized as callose stained with aniline blue, Pwl2 as green and HIPP43 magenta fluorescence respectively. Arrows are used to track Pwl2/HIPP43 co-localised mobile cytosolic puncta.

**Movie S2**

A laser confocal microscopy time-lapse video to show lack of co-localisation between AVR-PikE and HIPP43, 48 h after co-expression in *N. benthamiana* cells. PD are visualized as callose stained with aniline blue, AVR-PikE as green and HIPP43 magenta fluorescence respectively. Cyan arrows show co-localisation of aniline blue stain and HIPP43 as immobile signal at PD. Most of AVR-PikE expression occurred in the cytosol without HIPP43 co-localisation.

**Movie S3**

A laser confocal microscopy time-lapse video to show lack of co-localisation between MEP3 and HIPP43, 48 h after co-expression in *N. benthamiana* cells. PD are visualized as callose stained with aniline blue, MEP3 as green and HIPP43 magenta fluorescence respectively. Cyan arrows show co-localisation of aniline blue stain, MEP3 and HIPP43 as immobile signal at PD. Most of MEP3 expression occurred in the cytosol without HIPP43 co-localisation.

**Movie S4**

A laser confocal microscopy time-lapse video to show lack of co-localisation between a septuple mutant of Pwl2, Pwl2^SNDEYWY^ and HIPP43, 48 h after co-expression in *N. benthamiana* cells. PD are visualized as callose stained with aniline blue, Pwl2^SNDEYWY^ as green and HIPP43 magenta fluorescence respectively. Cyan arrows show co-localisation of aniline blue stain and HIPP43 as immobile signal at PD while expression of Pwl2^SNDEYWY^ remained in the cytosol without HIPP43 co-localisation

**Movie S5**

A laser confocal microscopy time-lapse video to show lack of co-localisation between free-YFP and HIPP43, 48 h after co-expression in *N. benthamiana* cells. laser confocal microscopy. PD are visualized as callose stained with aniline blue, free-YFP as green and HIPP43 magenta fluorescence respectively. Cyan arrows show co-localisation of aniline blue stain and HIPP43 as immobile signal at PD while expression of free-YFP remained in the cytosol without HIPP43 co-localisation.

**Table S1 Primers used in this study.**

**Supplemental Data 1. Blast genomes used for phylogenetic analysis.**

**Supplemental Data 2. Blast genomes used for copy number variation analysis.**

**Supplemental Data 3. IP-MS spectrum report.**

**Supplemental Data 4. IP-MS samples report with clusters.**

---

**Table S1.**
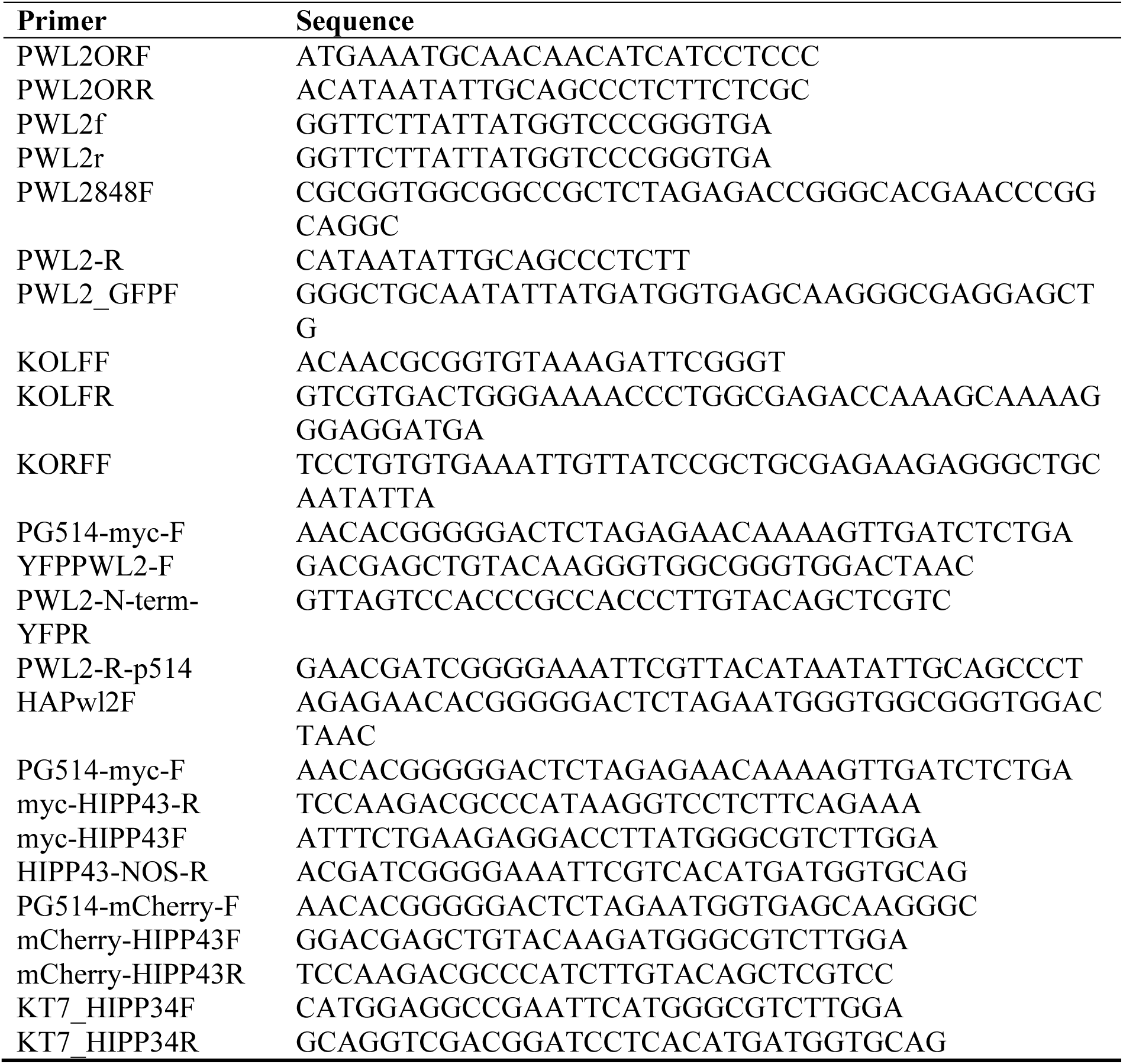
Primers used in this study.

